# Fast and exact single and double mutation-response scanning of proteins

**DOI:** 10.1101/2020.10.23.352955

**Authors:** Julian Echave

## Abstract

Studying the effect of perturbations on protein structure is a basic approach in protein research. Important problems, such as predicting pathological mutations and understanding patterns structural evolution, have been addressed by computational simulations based on modelling mutations as forces and predicting deformations using the Linear Response Approximation. In single mutation-response scanning simulations, a sensitivity matrix is obtained by averaging deformations over point mutations. In double mutation-response scanning simulations, a compensation matrix is obtained by minimizing deformations over pairs of mutations. These very useful simulation-based methods may be too slow to deal with large supra-molecular complexes, such as a ribosome or a virus capsid, or large number of proteins, such as the human proteome, which limits their applicability. To address this issue, I derived analytical closed formulas to calculate the sensitivity and compensation matrices directly, without simulations. Here, I present these derivations and show that the resulting analytical methods are much faster than their simulation counterparts, and that where the simulation methods are approximate, the analytical methods are exact by design.

## Introduction

Protein structure is considered to be fundamentally related to function. For this reason, insight into protein function can be gained by studying the structural deformations due to perturbations, such as those resulting from ligand binding or point mutations. This is at the basis of general experimental and theoretical approaches to study proteins. An experimental example is the powerful and increasingly popular Deep Mutational Scanning, which allows studying the effects of large numbers of mutations.^1,2^ Theoretically, the effect of small perturbations on protein structure has been studied using various approaches based on the Linear Response Approximation.^3–7^ Conformational deformations due to protein-ligand binding have been modelled using external forces applied locally to the superficial residues known or presumed to be involved in binding.^4,7^ This has proved useful for a number of applications such as predicting ligand-induced deformations,^4,7,8^ predicting ligand-binding sites related to known or desired deformations,^9,10^ and studying allosteric communication.^11–13^ Similar perturbation approaches focused on the structural response to mutations have been used for analysing pathological mutations^14,15^ and understanding patterns of protein evolution^6,14,16,17^

The previous simulation-based approaches may become too computationally costly to deal with very large systems, such as supra-molecular complexes, like a ribosome or a virus capsid. Also, scanning large sets of proteins, such as the whole human proteome to detect potential pathological mutations, may become impractical. The purpose of the present work to alleviate this problem by developing faster methods.

I focus on mutation-response scanning, considering two cases, single-mutation and double-mutation scanning. In simulation-based mutation-response scanning (`sMRS`), protein sites are scanned over, for each site many random mutations (modelled as forces) are introduced, the resulting deformations are calculated, and deformations are averaged over to obtain a sensitivity matrix **S**. In simulation-based double mutation-response scanning (`sDMRS`), pairs of sites are scanned over, random mutations are introduced at each sites of each pair, the resulting deformations are calculated, and the minimum deformations are used to calculate a compensation matrix **D**. As alternatives to these methods, I derive two analytical methods, `aMRS` and `aDMRS`, that allow, respectively, the calculation of **S** and **D** using closed-formed analytical formulas, without performing simulations. In the following sections, I describe the simulation methods, derive the analytical alternatives, and compare the speed and accuracy of simulation and analytical methods on a set of proteins of varying lengths.

## Methods

In the following sections, I derive the formalism of Mutation Response Scanning (`MRS`) and Double Mutation Response Scanning (`DMRS`). Key for these calculations, are the covariance matrix and the linear response approximation, explained next.

### Covariance matrix

At finite temperature the protein fluctuates, sampling an ensemble of conformations. Let a specific backbone conformation be specified by the position vector **r** = (*x*_1_, *y*_1_, *z*_1_, … *x*_*N*_, *y*_*N*_, *z*_*N*_)^*T*^, where (*x*_*i*_, *y*_*i*_, *z*_*i*_) are the Cartesian coordinates of site’s *i* alpha carbon, *N* is the number of sites, and super-index *T* denotes matrix or vector transposition. The native ensemble can be characterized by the *native structure*, **r**^0^ = ⟨**r**⟩, and by the *covariance matrix* :

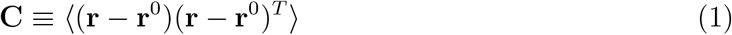

where ⟨· · ·⟩ is the average over conformations.

A protein’s conformational ensemble is determined by its energy landscape. For simplicity, in this work I use the Anisotropic Network Model (ANM).^18^ This model represents the protein as a network of amino acids connected by harmonic springs. Specifically, each residue is represented by single node placed at its *C*_*α*_, and pairs of nodes that are within a cut-off distance *R*_0_ are connected with springs of force-constant *k*. The ANM energy function is:

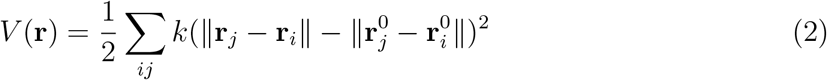

where **r**_*x*_ is the position vector of node *x*, **r**^0^_*x*_ its equilibrium position, *k* is the spring force constant, and the sum runs over all contacts *ij*. Regarding the model parameters, here I use a typical *R*_0_ = 12.5Å and *k* = 1.

Using (2), it is easy to obtain the covariance matrix. First, a second-order Taylor expansion of (2) leads to:

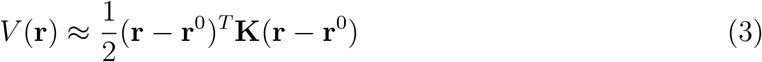

where 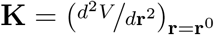 is the Hessian matrix. Then, assuming a Boltzmann distribution of conformations 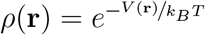 with *V* (**r**) given by (3), it follows that:

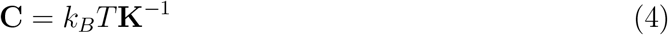

where *k*_*B*_ is Boltzmann’s constant, *T* the absolute temperature, and **K**^*−*1^ the Hessian’s pseudo-inverse. (**K** is not invertible because has 6 zero eigenvalues corresponding to rotations and translations.) Given a protein of known native structure **r**^0^, and parameters *R*_0_ and *k*, **K** is calculated differentiating (2), then **C** is obtained using (4).

### Linear Response Approximation

The covariance matrix determines the conformational shift that results from applying a force to one or more protein atoms. An arbitrary force can be represented by a vector **f** with one component for each of the coordinates that represent the protein’s conformation. For small **f**, the structural response can be calculated using the Linear Response Approximation (LRA):^4,6^

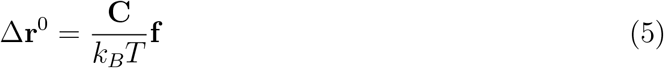

Equation (5) allows the prediction of the effect of any given force **f** with the sole knowledge of **C**.

### Mutation-Response Scanning

The aim of Mutation-Response Scanning (MRS) is to analyse how protein structure responds to point mutations. In the methods that I consider here, given a protein, mutations are modelled using forces, the resulting structural responses are calculated using the Linear Response Approximation, and these responses are averaged over mutations to calculate a sensitivity matrix **S** that quantifies the mutation-response patterns.

#### Mutations as forces

Point mutations can be modelled by forcing the contacts of the mutated site.^6^ Let *j* be the site to mutate, *C*(*j*) be the set of contacts of *j*, and *jl* the contact between sites *j* and *l*. Then, a mutation is modelled by applying a force

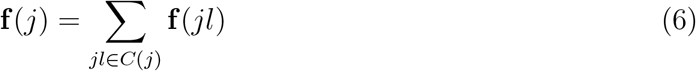

where the sum runs over contacts of *j* and **f** (*jl*) is the force applied to contact *jl*. Let *f* (*jl*) be a scalar and **e**_*jl*_ a unit vector directed from *j* to *l*. Then, **f** (*jl*) consists of a force *f* (*jl*)**e**_*jl*_ applied to *l*, plus a reaction force −*f* (*jl*)**e**_*jl*_ applied to *j*, and no force applied to other sites.

A random mutation of site *j* is modelled by picking independent random numbers *f* (*jl*) and building **f** (*jl*) and **f** (*j*) (Eq. 6). Following previous work,^6,16,17^ I use

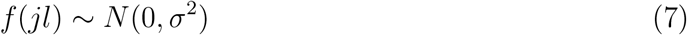

Thus, the contact forces are picked from independent identical normal distributions. (*σ*^2^ will affect the average size of the forces, which will have a mere scaling effect on mutation-response matrices.)

#### Sensitivity matrix, S

What is the effect on a site *i* of mutating a site *j*? Let us consider a random mutation at site *j*, represented by a force **f** (*j*). Then, from (5), the structural deformation due to this mutation is given by:

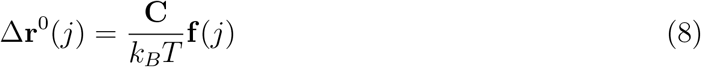

Δ**r** (*j*) can be written:

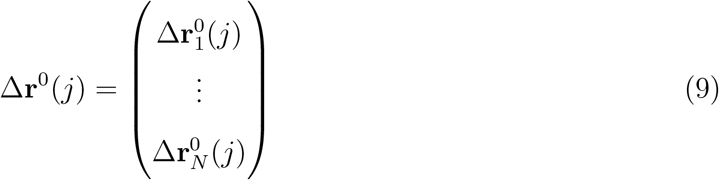

where Δ**r**_*i*_(*j*) is the 3 × 1 column vector that contains the change in Cartesian coordinates of site *i* due to mutation the given mutation **f** (*j*) applied to site j. The magnitude of the effect of the mutation on site’s *i* structure may be quantified by the Euclidean norm ||Δ**r**_*i*_(*j*)||^2^.

The *sensitivity matrix* **S** is the matrix with elements

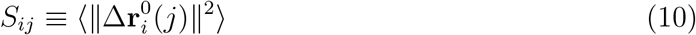

where *i* is the response site, *j* is the mutated site, and ⟨· · ·⟩ stands for averaging over random mutations of site *j*. *S*_*ij*_ represents the structural response of site *i* averaged over mutations at site *j*.

#### Simulation-based Mutation-Response Scanning

The sensitivity matrix **S** can be obtained using the simulation-based Mutation-Response Scanning method, `sMRS`. This numerical method proceeds as follows. For each site *j*, *M* random forces are generated following (6) and (7), the corresponding structural responses are calculated using (8), and averaged over mutations to calculate *S*_*ij*_, according to (10).

#### Analytical Mutation-Response Scanning

The key to the `sMRS` procedure is not the simulation itself, but the calculation of the sensitivity matrix **S**. In this section, I derive an alternative method: analytical Mutation-Response Scanning, `aMRS`.

Within the current model, mutating a site amounts to introducing independent forces to its contacts. Therefore, I first consider the deformation due to forcing a single contact. Let **f** (*jl*) be a force applied along contact *jl*, composed by a force *f* (*jl*)**e**_*jl*_ applied to *l* and a reaction force −*f* (*jl*)**e**_*jl*_ applied to *j*. Replacing **f** (*jl*) into (5) and using (9), we find

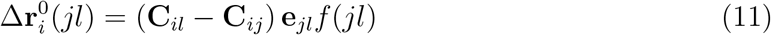

where 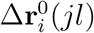 is the structural shift of site *i* due to the force applied to contact *jl* and **C**_*xy*_ is the 3 × 3 block of **C** corresponding to the covariance between sites *x* and *y*. Now, from (6), (8), and (9) we find

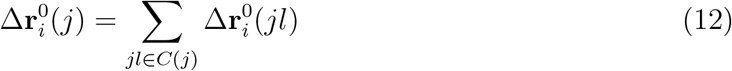

where 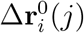 is the shift of *i* due to mutating site *j* and the sum runs over all contacts of *j*. Replacing (11) into (12), we find:

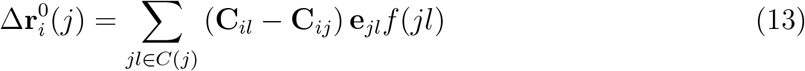

Replacing this equation into the definition of *S*_*ij*_, we get:

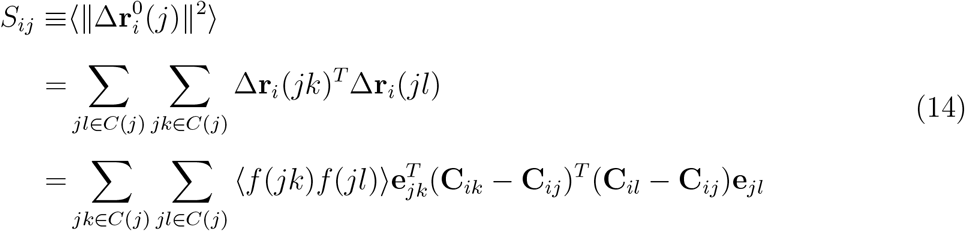

where ⟨· · ·⟩ stands for averaging over mutations at *j*. Since *f* (*jl*) ∼ *N* (0, *σ*^2^) are identically distributed independent random variables, (see **Mutations as forces**), it follows that

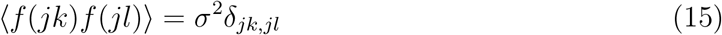

where *δ*_*xy*_ is the Kronecker delta, which is 1 for *x* = *y* and 0 otherwise. Replacing (15) into (14), we get:

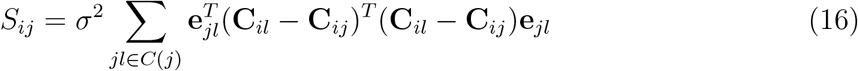

This equation allows the calculation of the sensitivity matrix without performing simulations. Given a protein, its structure **r**^0^ is used to calculate the unit vectors **e**_*kl*_ for all contacts, then these vectors and the protein’s covariance matrix **C** are used to calculate the sensitivity matrix elements using (16). This procedure constitutes the analytical mutation-response scanning method `aMRS`.

### Double Mutation-Response Scanning

The aim of Double Mutation-Response Scanning (DMRS) is to analyse how protein structure responds to pairs of point mutations. Just as for the MRS methods described above, the DMRS methods that I consider here model mutations using forces and calculate structural responses using the Linear Response Approximation. These responses are used to calculate a compensation matrix **D** that quantifies the degree of structural compensation between pairs of mutations.

#### Compensation matrix

To start, I define the compensation matrix that DMRS aims to calculate. Let Δ**r**^0^(*iμ*) be the structural response to a mutation *μ* at site *i*, and Δ**r**^0^(*jν*) be the structural response to a mutation *ν* at *j*. The deformation due to introducing both mutations is given by:

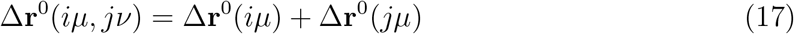

and the magnitude of this deformation is given by:

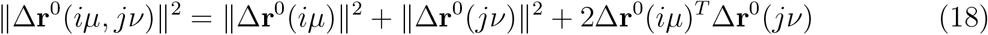

The first two terms are positive, but the third term may be positive or negative; when is negative, the mutations will compensate each other.

Given a first mutation *iμ*, the maximum compensation due to a second mutation at *j* is obtained when Δ**r**^0^(*iμ*)^*T*^ Δ**r**^0^(*jν*) is minimum; the degree of compensation is, therefore, 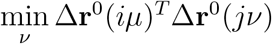. For mutations modelled as forces, this is equal to minus the maximum, because if a force maximizes the dot-product, the opposite force, which is as likely, minimizes it. Therefore, to keeps things positive, it is convenient to define the compensating power of *j* by 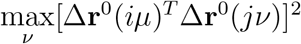.

To finish, I define a compensation matrix, **D**, with elements *D*_*ij*_ is given by:

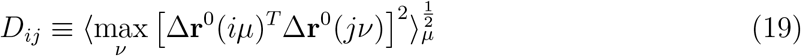

where ⟨· · ·⟩_*μ*_ is the average over *μ*. *D*_*ij*_ is a positive number that quantifies the degree to mutating *j* can compensate the structural effect of mutating *i*.

#### Simulation-based Double Mutation-Response Scanning

To obtain the compensation matrix, double-mutation simulations may be used. The method `sDMRS` (Simulation-Based Mutation-Response Scanning) proceeds as follows. First, for each site *k*, *M* forces **f** (*kμ*) are generated as described in **Mutations as forces**. Then, for each pair of sites (*i, j*), Δ**r**^0^(*iμ*) and Δ**r**^0^(*jν*) are obtained for all pairs of mutations (*iμ, jν*) using (5). Finally, compensation matrix elements *D*_*ij*_ are calculated using (19).

The previous `sDMRS` procedure demands adding another constraint to the forces. The value of Δ**r**^0^(*iμ*)^*T*^ Δ**r**^0^(*jν*) is proportional to the lengths of force vectors **f** (*iμ*) and **f** (*jμ*). Defined as described in **Mutations as forces**, the lengths of these vectors may become arbitrarily large, making the maximum in (19) infinite. To fix this, I apply the additional constraint:

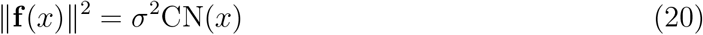

where *σ*^2^ is the parameter used to define contact forces (see Eq.7) and CN(*x*) is the number of contacts of site *x*. This specification ensures that it is not possible to have infinite forces.

#### Analytical DMRS

In this section, I derive the analytical Double Mutation-Response Scanning, `aDMRS`, method, an alternative way of calculating the compensation matrix without performing simulations.

Consider two mutations, at sites *i* and *j*, represented by forces **f** (*i*) and **f** (*j*), respectively. From (6) and (8), we find:

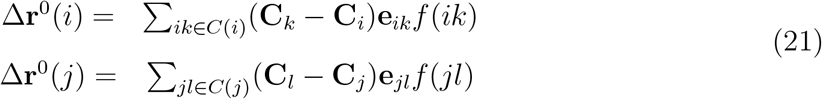

where Δ**r**^0^(*x*) is the protein’s deformation due to mutating site *x*, **C**_*x*_ is the 3*N* × 3 block of **C** with the 3 columns corresponding to site *x*, and *f* (*xy*) is the scalar force of contact *xy*.

Using this equation, we find:

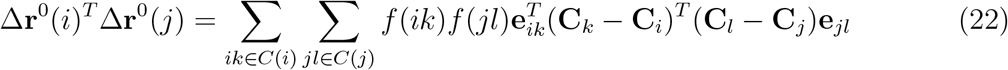

This equation can be written in matrix form:

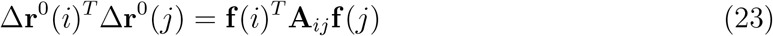

where **f** (*i*) is a column vector whose elements are the CN(*i*) contact forces *f* (*ik*), **f** (*j*) is the column vector with CN(*j*) elements *f* (*jl*), and **A**_*ij*_ is a matrix of size CN(*i*) × CN(*j*) with elements:

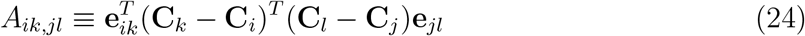

The maximum of [Δ**r**^0^(*i*)^*T*^ Δ**r**^0^(*j*)]^2^, subject to the constraint ||**f** (*j*)||^2^ = *σ*^2^CN(*j*) (Eq. 20). Using (23), it can be shown that:

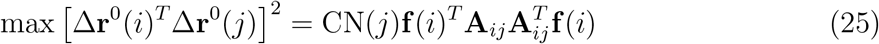

To finish, replacing (25) into (19), and using (15), we finally get:

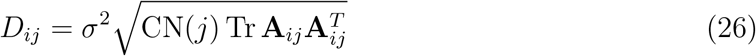

where Tr is the trace operator. This equation allows the calculation of the compensation matrix without performing simulations.

The method analytical Double Response Scanning, `aDMRS` proceeds as follows. Given a protein, its structure **r**^0^ is used to calculate the unit vectors **e**_*kl*_ for all contacts; then, these vectors and the protein’s covariance matrix **C** are used to calculate the matrices **A**_*ij*_ using (24); finally, (26) is used to calculate *D*_*ij*_.

## Results

To assess the analytical methods proposed in this work to their simulation-based counterparts, I applied all methods to the proteins of Table 1. The structure files for the calculations were obtained from the Protein Data Base for d2l8ma and d2acya, and from the Homstrad database for the other proteins.^19^

**Table 1:** Protein data set

| domain | family | class | $N$ |
| --- | --- | --- | --- |
| d1lcka1 | SH3 domain | All beta | 54 |
| d1ntxa | Snake venom toxins | Small | 60 |
| d1fxla2 | Canonical RNA-binding domain | Alfa & beta | 82 |
| d1bxva | Plastocyanine/Azurin-like | All beta | 91 |
| d2acya | Acyl-phosphatase-like | Alpha & beta | 98 |
| d1jiaa | Vertebrate Phospholipase A2 | All alpha | 122 |
| d1hmta | Fatty acid binding protein-like | All beta | 131 |
| d1a4fb | Globines | All alpha | 146 |
| d1mcta | Eukaryotic proteases | All beta | 223 |
| d2l8ma | Cytochrome P450 | All alpha | 405 |

Columns show, in order, protein domain id, family, and structural class according to the SCOP classification,<sup>20</sup> and protein length $N$ .

### Mutation-Response Scanning

In this section, the `sMRS` and `aMRS` methods are compared. Given a protein, an `sMRS` simulation consists in subjecting each of the protein sites *j* to *M* mutations, modelled as forces, calculating the resulting structural deformation of each other site *i*, and averaging these deformations over mutations to obtain a mutation-response matrix **S** (Eq. 10). `aMRS`, in contrast, calculates the mutational-response matrix without the need of simulating mutations, using the closed analytical expression Eq. 16. To assess these methods, I compare their computational speed and accuracy.

#### aMRS is much faster than sMRS

The computational speed of `sMRS` and `aMRS` is compared in Figure 1. An `sMRS` calculation using a typical number of *M* = 200 mutations per site is much slower than an `aMRS` calculation (Figure 1A). The computational cost, as measured by CPU time, scales with protein length as *N*^1.5^ for both `sMRS` and `aMRS`. As a result, *t*_sMRS_ increases linearly with *t*_aMRS_ with a slope that is the speedup of `aMRS` vs. `sMRS`; For the M = 200 case, *t*_sDMRS_ ≈ 126 × *t*_aDMRS_ (Figure 1B). Since `sDMRS` depends on *M* while `aDMRS` does not, the speedup depends on *M*. This dependence is linear: *t*_sMRS_*/t*_aMRS_ ∝ *M* (Figure 1C). Summarizing, the analytical method provides a speedup of the order of the number of mutations per site, which is typically in the hundreds; `aMRS` is much faster than `sMRS`.

**Figure 1:**
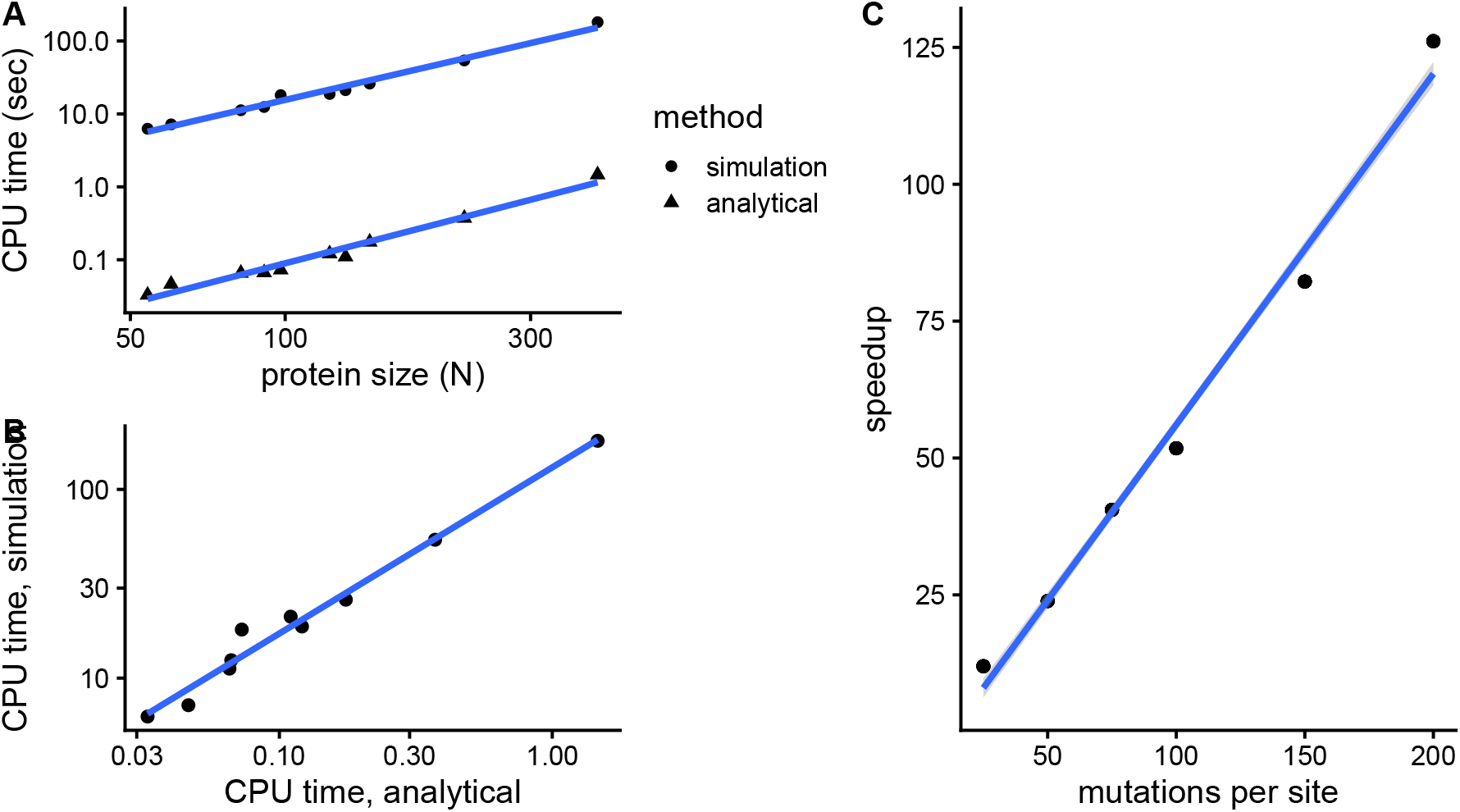
The analytical mutation-response method (aMRS) is much faster than the simulation method (sMRS). The simulation method calculates the sensitivity matrix averaging the structural response over random mutations applied to each of the protein sites (Eq. 10); the analytical method calculates the sensitivity matrix using a closed formula (Eq. 16). (A) CPU time vs. protein size the for `sMRS` with 200 mutations per site (simulation) and for `aMRS` (analytical). Both times scale with *N*^1.5^. (B) The CPU time of the simulation method increases linearly with the CPU time of the analytical method, with a speedup of 126: *t*_sMRS_ = 126 × *t*_aMRS_. (C) The speedup increases linearly with the number of mutations per site. Calculations performed on the proteins of Table 1 using the methods implemented in R, with base LAPACK and the optimized AtlasBLAS libraries for matrix operations, on an early-2018 MacBook Pro notebook (processor i7-8850H).

#### sMRS is very accurate, aMRS is exact

Regarding accuracy, in Figure 2, I compare simulated and analytical sensitivity matrices for the example case of Phospholipase A2 (SCOP id d1jiaa). By construction, the mutation-response matrix **S** calculated by `aMRS` is exact (Eq. 16). In contrast, `sMRS` is approximate and its accuracy will depend on the number of simulated mutations per site (*M*). Typically, *M* of *O*(10^2^) are used in mutation-response simulations (for some reason, *M* = 250 is quite common^11,21^). For 1djiaa, `sMRS` with *M* = 200 and `aMRS` lead to similar sensitivity matrices (Figure 2A and Figure 2B) and `sMRS` converges rapidly towards the exact `aMRS` matrix as *M* increases (Figure 2C). Thus, for Phospholipase A2, `sMRS` with a moderate number of mutations of *O*(10)-*O*(20^2^) produces accurate estimates of the exact `aMRS` matrix. Similar results are found for the other proteins of Table 1 (grey lines of Figure 2C, and `supplementary_info.pdf`).

**Figure 2:**
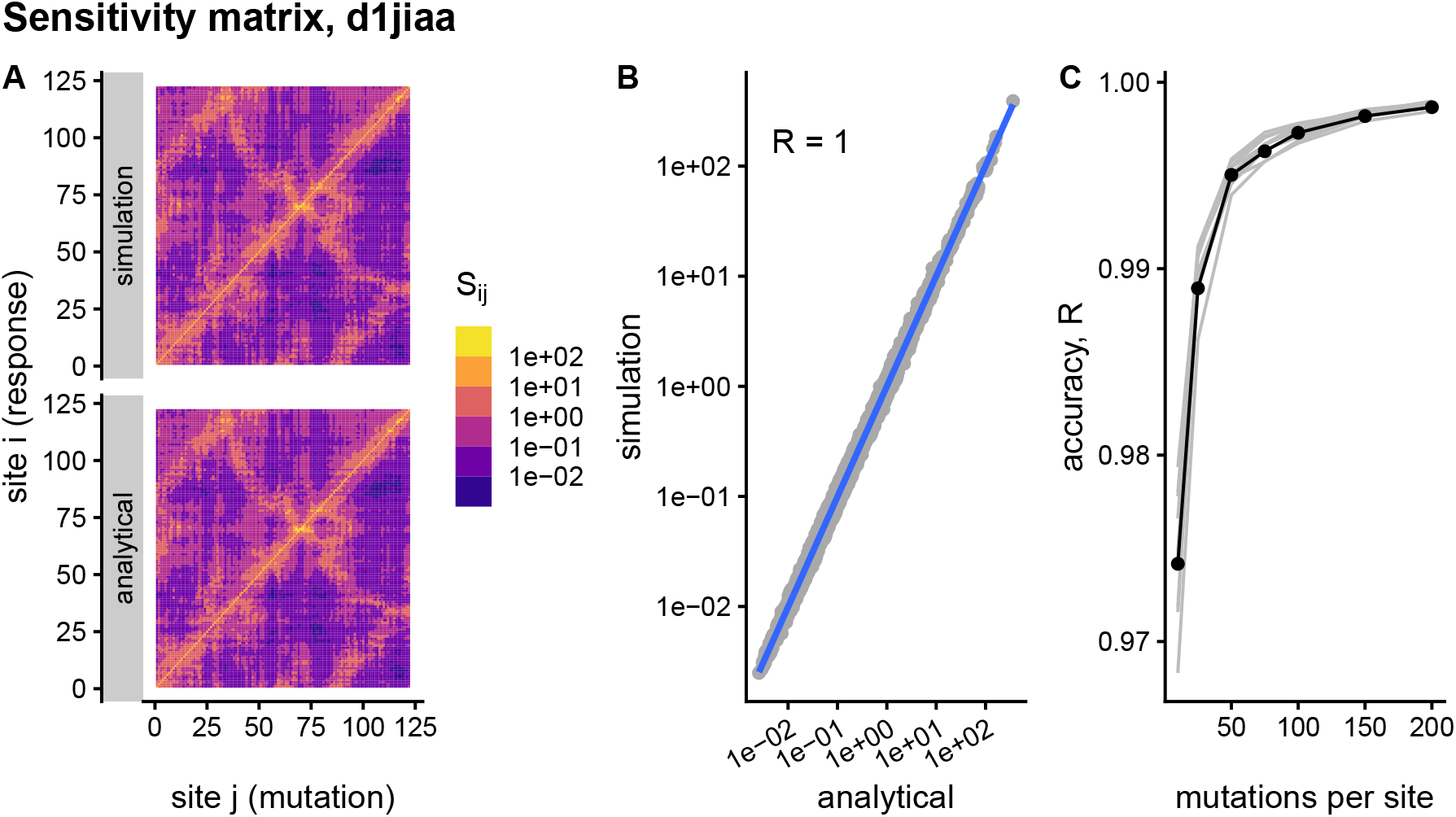
Comparison between sMRS and aMRS sensitivity matrices. Results shown for Phospholipase A2 (d1jiaa). The sensitivity matrix elements *S_ij_* measure the structural shift of site *i* averaged over mutations at site *j*. In `sMRS`, an approximate matrix is estimated by averaging over simulated mutations (Eq. 10); in `aMRS`, the exact matrix is obtained using an analytical closed formula (Eq. 16). (A) `sMRS` response matrix obtained by averaging over 200 mutations (simulation) compared with the exact `aMRS` matrix (analytical). (B) `sMRS` vs. `aMRS` matrix elements; points lie close to the *y* = *x* diagonal. (C) As *M* increases, `sMRS` converges rapidly towards high correlations with the exact `aMRS` results; the d1jiaa case is shown with black lines and points, and the other 9 proteins studied are shown using grey lines. Matrices are normalized so that their average is 1. Logarithmic scale is used in A and B and *R* is the Pearson correlation coefficient between the log-transformed `sMRS` and `aMRS` matrices.

Averaging the sensitivity matrix over rows or columns we obtain one-dimensional site-dependent profiles. Column *j* of **S** is the profile of responses of sites *i* to mutating *j*, and, therefore, it measures the influence of *j* over other sites. Therefore, averaging over *i*, we obtain the mean *influence profile* with elements 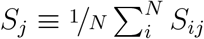. On the other hand, a row *i* of **S** is the profile of responses of *i* to mutations at sites *j*, which measures the *sensitivity* if *i*. Accordingly, averaging over *j* we obtain the mean *sensitivity profile* with elements 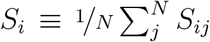. In general, **S** is not symmetric and, therefore, sensitivity and influence profiles are different.

To further assess accuracy, Figure 3 compares `sMRS` and `aMRS` profiles for Phospholipase A2 (d1jiaa). Comparing influence profiles, we see that `sMRS` with *M* = 200 gives accurate estimates of the exact `aMRS` profiles (Figure 3A and Figure 3B) and that `sMRS` converges rapidly towards the exact `aMRS` influence profiles as *M* increases (Figure 2C). Similarly, sensitivity profiles are also very accurately estimated by `sMRS` with *M* = 200 (Figures 3D and 3E) and `sMRS` converges rapidly towards the exact `aMRS` profiles as *M* increases (Figure 2F). Note that `sMRS` influence profiles are less accurate than `sMRS` sensitivity profiles (compare Figure 3B with Figure 3E and Figure 3C with Figure 3F). However,with a typical number of *M* = *O*(10^2^) mutations, both profiles are very well converged. Thus, `sMRS` with *M* of *O*(10^2^) provides accurate estimates of the exact `aMRS` sensitivity and influence profiles for Phospholipase A2. Similar results are found for the other proteins of the Table 1 (argy lines of Figure 2C, and `supplementary_info.pdf`).

**Figure 3:**
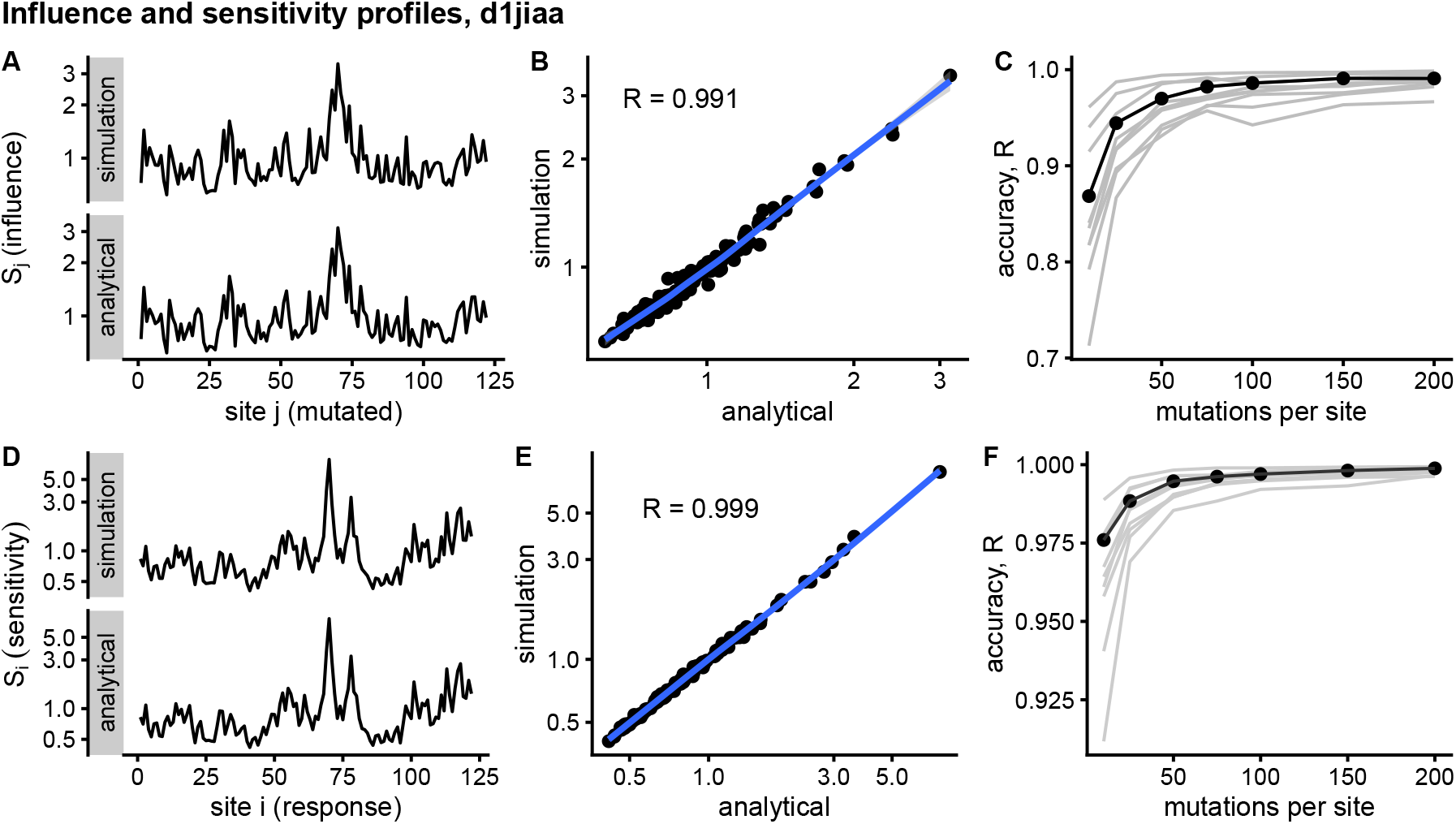
Comparison of sMRS and aMRS marginal profiles. Results shown for Phospholipase A2 (d1jiaa). The influence profile is the average of the sensitivity matrix over rows; element *S_j_* measures the influence of mutations at site *j* to deform the protein. The sensitivity profile is the average of the response matrix over columns; element *S_i_* measures the average sensitivity of site *i*. The simulation-based and analytical methods, `sMRS` and `aMRS`, are described in the caption of Figure 2. (A) Comparison of `sMRS` and `aMRS` influence profiles; (B) scatter plot of the approximate `sMRS` vs. exact `aMRS` *S_j_* values of (A); (C) convergence of the `sMRS` *S_j_* profiles towards the exact `aMRS` profile. (D) Comparison of `sMRS` and `aMRS` sensitivity profiles; (E) scatter plot of the approximate `sMRS` vs. exact `aMRS` *S_i_* values of (D); (F) convergence of the `sMRS` *S_i_* profiles towards the exact `aMRS` profile. In (C) and (F), the d1jiaa case is shown with black lines and points, and the other 9 proteins studied are shown using grey lines. Profiles calculated using the normalized matrix (matrix average is 1). Profile elements are shown in logarithmic scale and *R* is the Pearson correlation coefficient between the log-transformed `sMRS` and `aMRS` profiles.

### Double mutation-response scanning

This section compares the two double mutation-response scanning methods, `sDMRS` and `aDMRS`. These methods are alternative ways of calculating a compensation matrix **D** with elements *D_ij_* that measure the compensation between structural deformations due to a first mutation mutation at *i* and a second mutation at *j* (Eq.19). The simulation method `sDMRS` obtains this matrix numerically by maximizing the compensation over pairs of mutations at each pair of sites. The analytical method `aDMRS` calculates the compensation values using a closed formula (Eq. 26).

#### aDMRS is much faster than sDMRS

The computational speed of `sDMRS` and `aDMRS` is compared in Figure 4. `sDMRS` with *M* = 200 mutations per site is much slower than `aDMRS` (Figure 4A). The computational cost, as measured by CPU time, scales with protein length as *N*^3^ for both `sDMRS` and `aDMRS`. As a result, *t*_sDMRS_ increases linearly with *t*_aDMRS_ with a slope that is the speedup of `aDMRS` vs. `sDMRS`. For the 200-mutations-per-site case, *t*_sDMRS_ ≈ 137×*t*_aDMRS_ (Figure 4B). Since the computational cost of `sDMRS` increases with *M* while the cost of `aDMRS` does not, the speedup depends on *M*. This dependence is non-linear (Figure 4C); as *M* increases, the cost of generating the mutations increases linearly with *M*, while performing the average and maximization needed to calculate the compensation matrix scales as *M*^2^. In summary, for large *M* the analytical method provides a speedup of *O*(*M*^2^), making `aMRS` much faster than `sMRS`.

**Figure 4:**
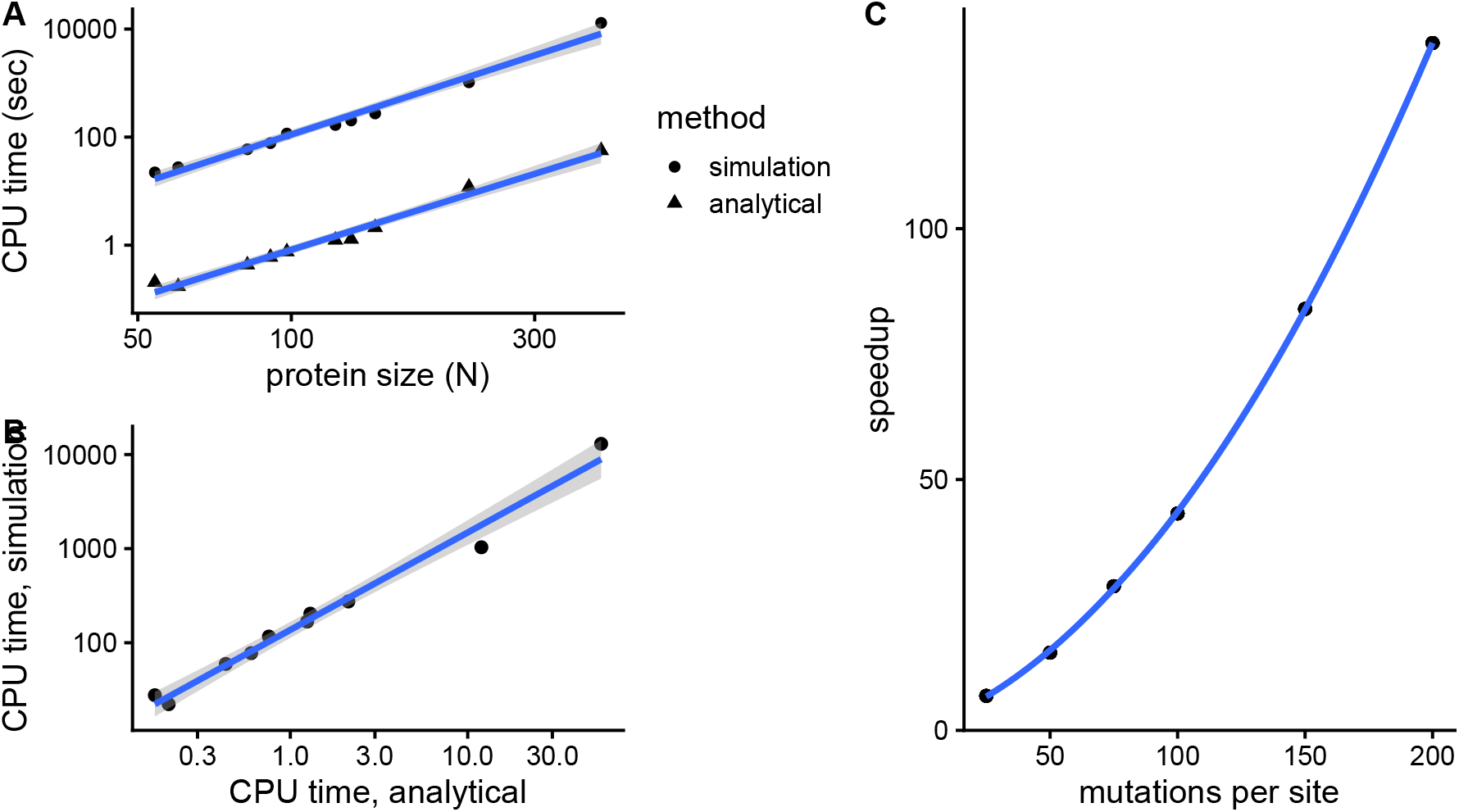
The analytical double mutation-response scanning method (aDMRS) is much faster than the simulation method (aDMRS). `sDMRS` calculates a compensation matrix by maximizing the structural compensation over numerically generated random mutations, modelled as forces applied to pairs of sites (Eq. 19); the analytical method calculates this matrix using a closed formula (Eq. 26). (A) CPU time vs. protein size the for `sDMRS` with 200 mutations per site (simulation) and for `aDMRS` (analytical). Both times scale with *N*^3^. (B) The CPU time of the simulation method increases linearly with the CPU time of the analytical method, with a speedup of 137: *t*_sMRS_ = 137 × *t*_aMRS_. (C) The speedup increases non-linearly with the number of mutations per site, tending towards *O*(*M*^2^) for large *M*. Calculations performed on the proteins of Table 1 using the methods implemented in R, with base LAPACK and the optimized AtlasBLAS libraries for matrix operations, on an early-2018 MacBook Pro notebook (processor i7-8850H).

#### sDMRS is moderately accurate, aDMRS is exact

Regarding accuracy, in Figure 5, I compare simulated and analytical response matrices for the example case of Phospholipase A2 (SCOP id d1jiaa). By construction, `aDMRS` is exact, while `sDMRS` is approximate. The accuracy of `sDMRS` will depend on the number of mutations per site (*M*) over which compensation is maximized. The compensation matrices obtained with `sDMRS` with *M* = 200 looks similar to the exact `aDMRS` matrix. (Figure 5A). A scatter plot `sDMRS` vs. `aDMRS` matrix elements shows good correlation, but a visible scattering of points around the linear fit (Figure 5C). The similarity can be measured by the correlation coefficient, which which in this case is *R* = 0.92. The accuracy of `sDMRS` improves very slowly as *M* increases (Figure 5C). Thus, for Phospholipase A2, `sDMRS` with *O*(10^2^) mutations per site produces a compensation matrix that is in moderately good agreement with the exact `aDMRS` matrix. Results are similar for other proteins (see grey lines of Figure 5C and `supplementary_info.pdf`).

**Figure 5:**
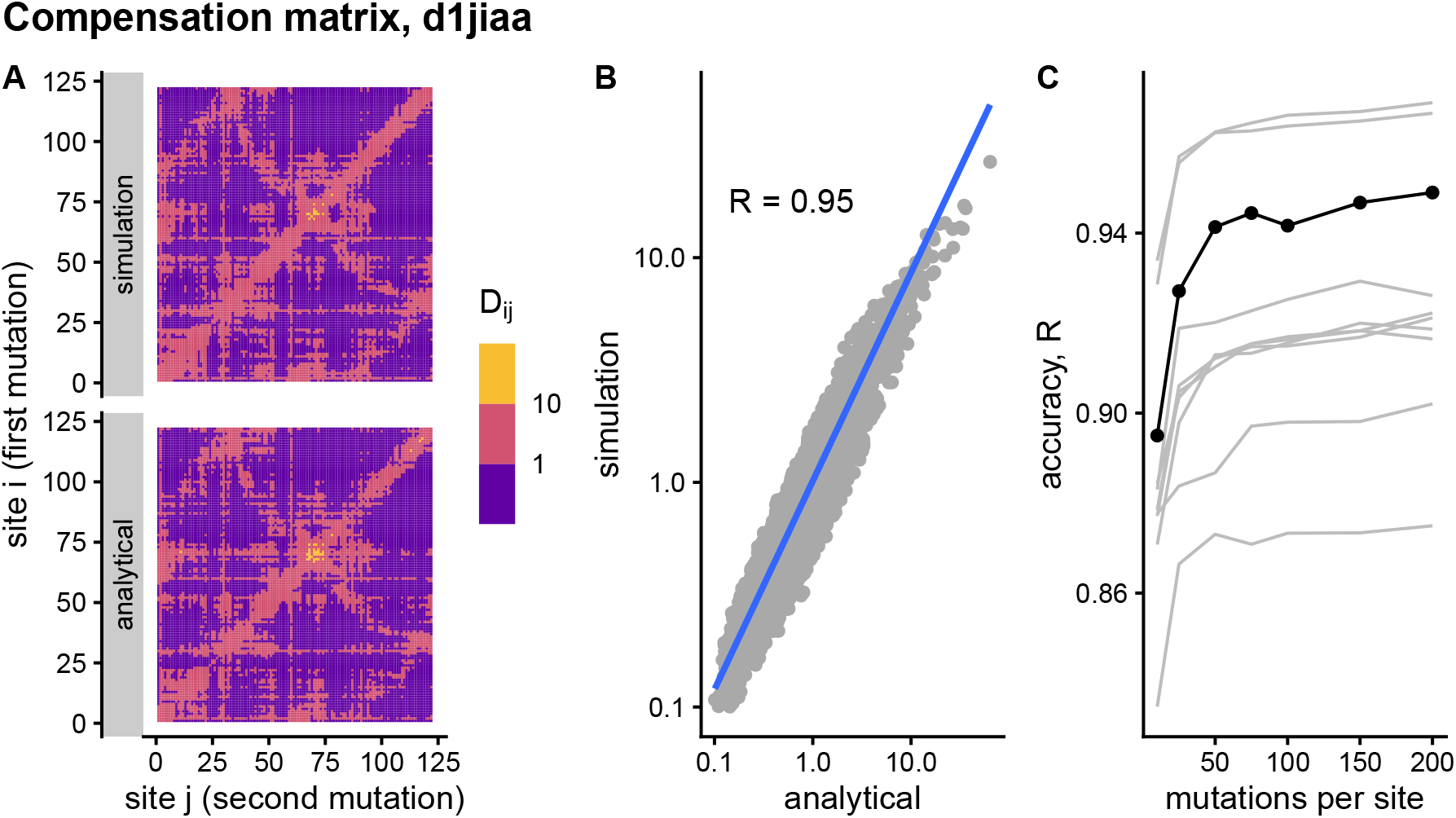
Comparison of sDMRS and aDMRS compensation matrices. Results shown for Phospholipase A2 (d1jiaa). The compensation matrix elements *D*_*ij*_ measure the maximum compensation of the structural deformation due to a mutation at site *i* afforded by a second mutation at *j*. In `sDMRS`, *D*_*ij*_ are obtained by maximizing structural compensation over simulated pairs of mutations (Eq. 19); in `aDMRS`, the exact *D_ij_* are obtained using an analytical closed formula (Eq. 26). (A) `sDMRS` compensation matrix, obtained by maximizing over 200 mutations per site (simulation) compared with the `aDMRS` matrix (analytical). (B) `sDMRS` vs. `aDMRS` matrix elements. (C) Convergence of the `sDMRS` matrix towards the exact `aDMRS` matrix with increasing number of mutations per site. In (C) the d1jiaa case is shown with black lines and points, and the other 9 proteins studied are shown using grey lines. `sMRS` converges very slowly towards the exact `aDMRS` results. *D*_*ij*_ are normalized so that their average is 1, logarithmic scales are used in (A) and (B), and *R* is Pearson’s correlation coefficient between log-transformed `sMRS` and `aMRS` matrix elements.

Using the compensation matrix we can obtain compensation profiles. Averaging **D** over rows, we obtain a *D*_*j*_ profile that measures the average compensation power of sites *j*. Averaging over columns we obtain a *D*_*i*_ profile that measures how likely to be compensated are mutations at *i*.

Figure 6 compares `sMRS` and `aMRS` profiles for Phospholipase A2 (d1jiaa). Profiles obtained using `sMRS` with *M* = 200 are similar to `aDMRS` profiles (Figure 6A and Figure 6D). The similarity is not very high, however: points are quite scattered around the linear fit in `sDMRS` vs. `aDMRS` plots (Figure 6B and Figure 6E). Convergence of `sDMRS` towards `aDMRS` is very slow (Figure 6C and Figure 6F). The case of d1jiaa is not the worst. For other proteins, the correlation between `sDMRS` and `aDMRS` profiles varies between 0.58 and 0.98 (see grey lines of Figure 6C and Figure 6F, and `supplementary_info.pdf`). In summary, `sDMRS` with *M* of *O*(10^2^) provides compensation profiles that are in poor to good agreement with exact `aDMRS` profiles, depending on the protein.

**Figure 6:**
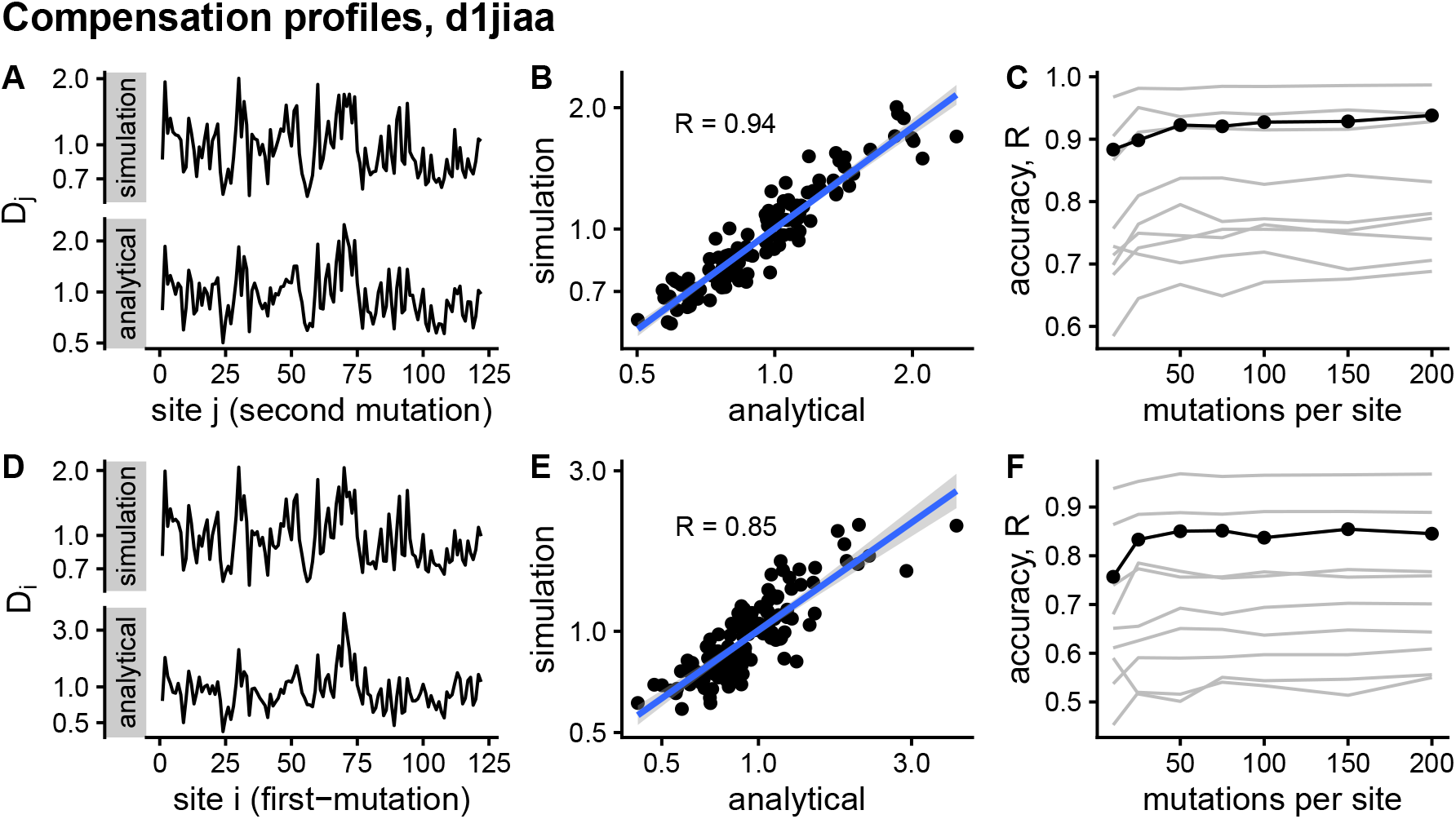
Comparison of sDMRS and aDMRS marginal profiles. Results shown for Phospholipase A2 (d1jiaa). Two marginal profiles are considered. The *D*_*j*_ profile is the average of the compensation matrix over rows; element *D*_*j*_ measures the ability of *j* to compensate mutations at other sites. The *D*_*i*_ profile is the average of the compensation matrix over columns; element *D*_*i*_ measures the the degree to which a mutation at *i* can be compensated by mutations elsewhere. The simulation-based and analytical methods, `sDMRS` and `aDMRS`, are described in the caption of Figure 5. (A) Comparison of `sDMRS` and `aDMRS` *D*_*j*_ profiles; (B) scatter plot of the approximate `sDMRS` vs. exact `aDMRS` *D*_*j*_ values of (A); (C) convergence of the `sDMRS` *D*_*j*_ profiles towards the exact `aDMRS` profile. (D) Comparison of `sDMRS` and `aDMRS` *D*_*i*_ profiles; (E) scatter plot of the approximate `sDMRS` vs. exact `aDMRS` *D*_*i*_ values of (D); convergence of the `sDMRS` *D*_*i*_ profiles towards the exact `aDMRS` profile. In (C) and (F), the d1jiaa case is shown with black lines and points, and the other 9 proteins studied are shown using grey lines. Profiles were calculated with normalized matrices (matrix average is 1), they are in logarithmic scale, and *R* is the Pearson correlation coefficient between the log-transformed `sDMRS` and `aDMRS` profiles.

## Discussion

In the previous sections I have presented and assessed two mutation-response scanning methods, `aMRS` and `aDMRS`, which are analytical alternatives to the simulation methods `sMRS` and `sDMRS`, respectively. These methods have two advantages over the simulation methods: speed and accuracy.

First: the analytical methods are much faster than the simulation methods. For a typical case of *M* = 200 mutations per site, `aMRS` is 126× faster than `sMRS` and `aDMRS` is 137× faster than `sDMRS`. While the computational cost of `sMRS` is relatively modest and increases rather slowly in proportion to *N*^1.5^*M*, `sDMRS` is much more computationally expensive and cost rises steeply in proportion to *N*^3^*M*^2^. The speedup of analytical methods is of *O*(*M*) for single-mutation scans and *O*(*M*^2^) for double-mutation scans. Therefore, in both cases the speedup is important, but it may be most useful for double-mutation scans of large proteins. For instance, for the 405-sites-long Cytochrome P450, calculating **S** is takes 3 CPU minutes using simulations vs. 1.5 seconds of the analytical method. Calculating **D**, on the other hand, takes 3.6 hours using simulations vs. 1 minute using the analytical method (Table S1 and Table S2).

Second: while the simulation methods are approximate, the analytical methods are exact. `aMRS` allows the exact calculation of the sensitivity matrix **S** (Eq. 16) and `aDMRS` the exact calculation of the compensation matrix **D** (Eq. 26). In contrast, the accuracy of simulation methods depends on the number of mutations per site *M*. With a typical *M* = *O*(10^2^), `sMRS` provides very accurate approximations to **S** and its marginal profiles (Table S1). On the other hand, `sDMRS` converges very slowly, so that even with *M* = *O*(10^2^) `sDMRS` compensation matrices and profiles are poorly converged (Table S2). The reason why `sDMRS` converges more slowly than `sMRS` is that it is more difficult to find extreme values (calculation of the compensation matrix involves maximization over pairs of mutations) than averages (sensitivity matrix elements are averages over mutations). Therefore, from the point of view of accuracy, the analytical option is better both for single-point and double-point scans, but it is most important for the latter.

In conclusion, the analytical methods presented here should be the methods of choice to calculate mutation-response matrices and compensation matrices, because they are much faster and exact. The speedup afforded by these methods, would be especially useful to analyse otherwise intractable large proteins, protein complexes, and large protein databases. These methods should be useful for a wide range of potential applications, such as predicting evolutionary divergence of protein structures,^16,17^ detecting and interpreting pathological mutations,^14,21,22^ and detecting compensating mutations and rescue sites.^15^

To finish, I mention two possible lines of further development. A first line is to derive analytical expressions for the deformations due to external forces applied to single sites, as in Perturbation-Response Scanning (PRS)^7,11^ and the Double Force Scanning (DFS).^15^ This will be useful for applications related to ligand-binding induced deformations.^9,11^ Beyond deformations, a second line of development is to derive analytical alternatives to simulation-based methods that calculate effects of mutations on protein motions.^23,23–26^ This would be important for studies of the role of protein dynamics in function and evolution.^26–31^

## Acknowledgement

This work was supported by Consejo Nacional de Investigaciones Científicas y Técnicas (grant number PIP 112 201501 00385 CO) and by Agencia Nacional de Promoción Científica y Tecnológica (grant number PICT-2016-4209).

## Supporting Information Available

Supplementary information file: `supplementary_info.pdf`

Data: https://doi.org/10.5281/zenodo.4122623

Code: https://doi.org/10.5281/zenodo.4123149

## Notes

https://doi.org/10.5281/zenodo.4122623

### Competing Interest Statement

The authors have declared no competing interest.

## References

(1) Fowler, D. M.; Fields, S. Deep mutational scanning: A new style of protein science. Nat. Methods 2014, 11, 801–807.

(2) Livesey, B. J.; Marsh, J. A. Using deep mutational scanning to benchmark variant effect predictors and identify disease mutations. Mol. Syst. Biol. 2020, 16, 1–12.

(3) Yilmaz, L. S.; Atilgan, A. R. Identifying the adaptive mechanism in globular proteins: Fluctuations in densely packed regions manipulate flexible parts. J. Chem. Phys. 2000, 113, 4454–4464.

(4) Ikeguchi, M.; Ueno, J.; Sato, M.; Kidera, A. Protein structural change upon ligand binding: Linear response theory. Phys. Rev. Lett. 2005, 94, 1–4.

(5) Zheng, W.; Brooks, B. R. Normal-modes-based prediction of protein conformational changes guided by distance constraints. Biophys. J. 2005, 88, 3109–3117.

(6) Echave, J. Evolutionary divergence of protein structure: The linearly forced elastic network model. Chem. Phys. Lett. 2008, 457, 413–416.

(7) Atilgan, C.; Atilgan, A. A. R. Perturbation-response scanning reveals ligand entry-exit mechanisms of ferric binding protein. PLoS Comput. Biol. 2009, 5.

(8) Tamura, K.; Hayashi, S. Linear Response Path Following: A Molecular Dynamics Method To Simulate Global Conformational Changes of Protein upon Ligand Binding. J. Chem. Theory Comput. 2015, 11, 2900–17.

(9) Atilgan, C.; Gerek, Z. N.; Ozkan, S. B.; Atilgan, A. R. Manipulation of conformational change in proteins by single-residue perturbations. Biophys. J. 2010, 99, 933–943.

(10) Jalalypour, F.; Sensoy, O.; Atilgan, C.; Atilgan, C. Perturb-Scan-Pull: A Novel Method Facilitating Conformational Transitions in Proteins. J. Chem. Theory Comput. 2020, 16, 3842–3855.

(11) General, I. J.; Liu, Y.; Blackburn, M. E.; Mao, W.; Gierasch, L. M.; Bahar, I. ATPase Subdomain IA Is a Mediator of Interdomain Allostery in Hsp70 Molecular Chaperones. PLoS Comput. Biol. 2014, 10.

(12) Alfayate, A.; Caceres, C. R.; Dos Santos, H. G. H.; Bastolla, U. Predicted dynamical couplings of protein residues characterize catalysis, transport and allostery. Bioinformatics 2019, 35, 4971–4978.

(13) Lake, P. T.; Davidson, R. B.; Klem, H.; Hocky, G. M.; McCullagh, M. Residue-Level Allostery Propagates through the Effective Coarse-Grained Hessian. J. Chem. Theory Comput. 2020, 16, 3385–3395.

(14) Nevin Gerek, Z.; Kumar, S.; Banu Ozkan, S. Structural dynamics flexibility informs function and evolution at a proteome scale. Evol. Appl. 2013, 6, 423–433.

(15) Tiberti, M.; Pandini, A.; Fraternali, F.; Fornili, A. In silico identification of rescue sites by double force scanning. Bioinformatics 2018, 34, 207–214.

(16) Echave, J.; Fernández, F. M. A perturbative view of protein structural variation. Proteins Struct. Funct. Bioinforma. 2010, 78, 173–180.

(17) Marcos, M. L.; Echave, J. The variation among sites of protein structure divergence is shaped by mutation and scaled by selection. Curr. Res. Struct. Biol. 2020, 2, 156–163.

(18) Atilgan, A. R.; Durell, S. R.; Jernigan, R. L.; Demirel, M. C.; Keskin, O.; Bahar, I. Anisotropy of fluctuation dynamics of proteins with an elastic network model. Biophys. J. 2001, 80, 505–515.

(19) Stebbings, L. A.; Mizuguchi, K. HOMSTRAD: Recent developments of the Homologous Protein Structure Alignment Database. Nucleic Acids Res. 2004, 32, 203D–207.

(20) Murzin, A. G.; Brenner, S. E.; Hubbard, T.; Chothia, C. SCOP: A structural classification of proteins database for the investigation of sequences and structures. J. Mol. Biol. 1995, 247, 536–540.

(21) Verkhivker, G. M. Biophysical simulations and structure-based modeling of residue interaction networks in the tumor suppressor proteins reveal functional role of cancer mutation hotspots in molecular communication. Biochim. Biophys. Acta - Gen. Subj. 2019, 1863, 210–225.

(22) Raimondi, D.; Orlando, G.; Tabaro, F.; Lenaerts, T.; Rooman, M.; Moreau, Y.; Vranken, W. F. Large-scale in-silico statistical mutagenesis analysis sheds light on the deleteriousness landscape of the human proteome. Sci. Rep. 2018, 8, 1–11.

(23) Hamacher, K. Relating sequence evolution of HIV1-protease to its underlying molecular mechanics. Gene 2008, 422, 30–36.

(24) Zheng, W.; Tekpinar, M. Large-scale evaluation of dynamically important residues in proteins predicted by the perturbation analysis of a coarse-grained elastic model. BMC Struct. Biol. 2009, 9, 45.

(25) Zheng, W.; Thirumalai, D. Coupling between normal modes drives protein conformational dynamics: Illustrations using allosteric transitions in myosin II. Biophys. J. 2009, 96, 2128–2137.

(26) Echave, J. Why are the low-energy protein normal modes evolutionarily conserved? Pure Appl. Chem. 2012, 84, 1931–1937.

(27) Micheletti, C. Comparing proteins by their internal dynamics: Exploring structure-function relationships beyond static structural alignments. Phys. Life Rev. 2013, 10, 1–26.

(28) Ponzoni, L.; Bahar, I. Structural dynamics is a determinant of the functional significance of missense variants. Proc. Natl. Acad. Sci. U. S. A. 2018, 115, 4164–4169.

(29) Zhang, S.; Li, H.; Krieger, J. M.; Bahar, I.; Ozkan, B. Shared Signature Dynamics Tempered by Local Fluctuations Enables Fold Adaptability and Specificity. Mol. Biol. Evol. 2019, 36, 2053–2068.

(30) Zhang, P. F.; Su, J. G. Identification of key sites controlling protein functional motions by using elastic network model combined with internal coordinates. J. Chem. Phys. 2019, 151.

(31) Wingert, B.; Krieger, J.; Li, H.; Bahar, I. Adaptability and specificity: how do proteins balance opposing needs to achieve function? Curr. Opin. Struct. Biol. 2021, 67, 25–32.

