## Supplementary info for "Fast and exact single and double mutation-response scanning of proteins"

Julian Echave*

*Instituto de Ciencias Físicas, Escuela de Ciencia y Tecnología, Universidad Nacional de San Martín, Martín de Irigoyen 3100, 1650 San Martín, Buenos Aires, Argentina*

### Supplementary Tables

Table S1: Mutation-response scanning, simulation ($M = 200$) vs. analytical

| protein | $N$ | simulation CPU time | analytical CPU time | speedup | accuracy ($R$) |
|---|---|---|---|---|---|
| d1lcka1 | 54 | 6.26 | 0.03 | 126 | 1.00 |
| d1ntxa | 60 | 7.18 | 0.05 | 126 | 1.00 |
| d1fxla2 | 82 | 11.22 | 0.07 | 126 | 1.00 |
| d1bxva | 91 | 12.44 | 0.07 | 126 | 1.00 |
| d2acya | 98 | 18.08 | 0.07 | 126 | 1.00 |
| d1jiaa | 122 | 18.77 | 0.12 | 126 | 1.00 |
| d1hmta | 131 | 21.16 | 0.11 | 126 | 1.00 |
| d1a4fb | 146 | 26.02 | 0.18 | 126 | 1.00 |
| d1mcta | 223 | 54.08 | 0.37 | 126 | 1.00 |
| d2l8ma | 405 | 180.91 | 1.47 | 126 | 1.00 |

Table S2: Double mutation-response scanning, simulation ($M = 200$) vs. analytical

| protein | $N$ | simulation CPU time | analytical CPU time | speedup | accuracy ($R$) |
|---|---|---|---|---|---|
| d1lcka1 | 54 | 22.24 | 0.21 | 137 | 0.87 |
| d1ntxa | 60 | 27.70 | 0.17 | 137 | 0.97 |
| d1fxla2 | 82 | 59.58 | 0.43 | 137 | 0.93 |
| d1bxva | 91 | 77.46 | 0.60 | 137 | 0.92 |
| d2acya | 98 | 116.38 | 0.76 | 137 | 0.90 |
| d1jiaa | 122 | 167.89 | 1.24 | 137 | 0.95 |
| d1hmta | 131 | 203.65 | 1.30 | 137 | 0.97 |
| d1a4fb | 146 | 274.29 | 2.13 | 137 | 0.92 |
| d1mcta | 223 | 1034.36 | 11.95 | 137 | 0.92 |
| d2l8ma | 405 | 12995.91 | 56.53 | 137 | 0.92 |

### Supplementary Figures

The following pages contains figures similar to those of the main document, for other proteins.

See captions of main figures for details.

**Sensitivity matrix, d1lcka1**

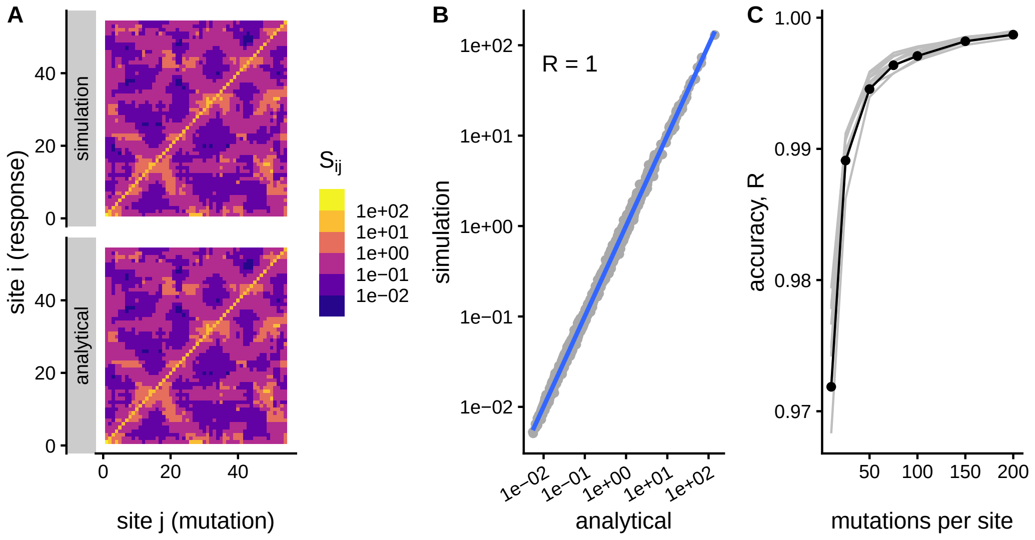


**Influence and sensitivity profiles, d1lcka1**

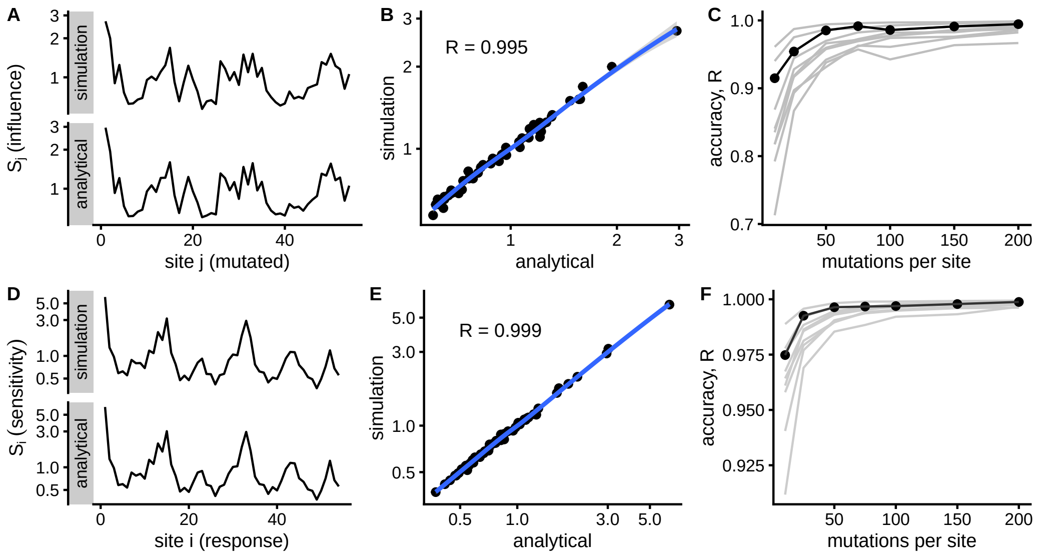

#### Compensation matrix, d1lcka1

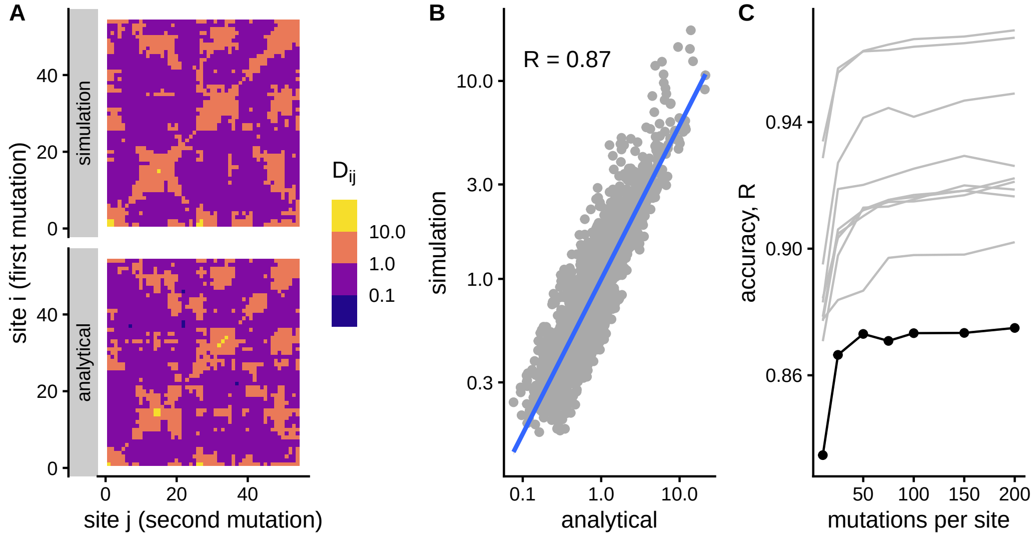


#### Compensation profiles, d1lcka1

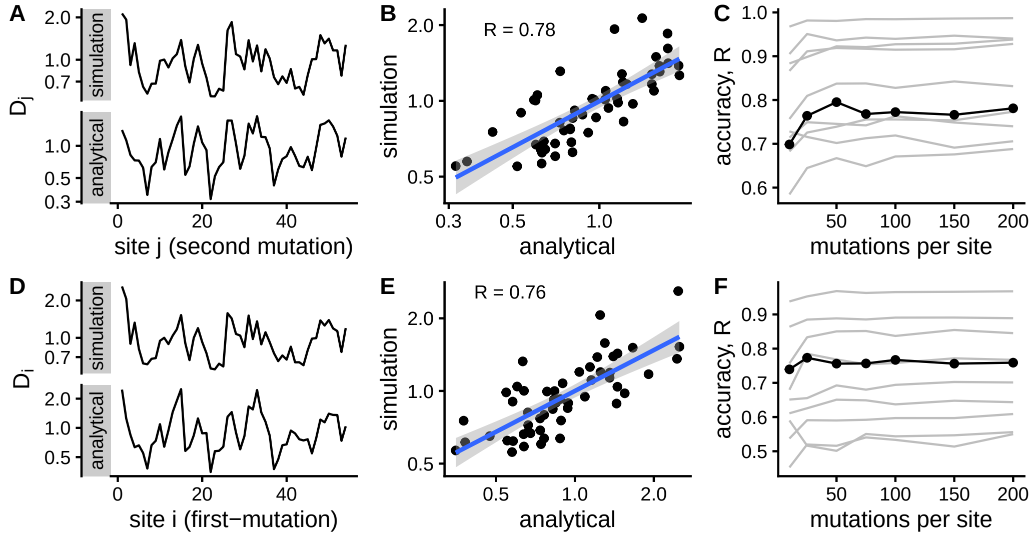

**Sensitivity matrix, d1ntxa**

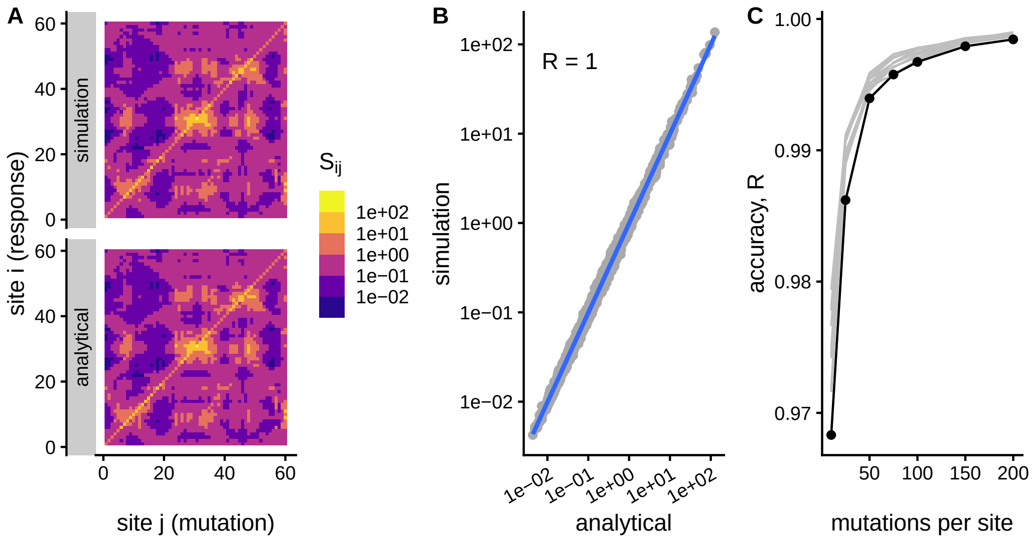


**Influence and sensitivity profiles, d1ntxa**

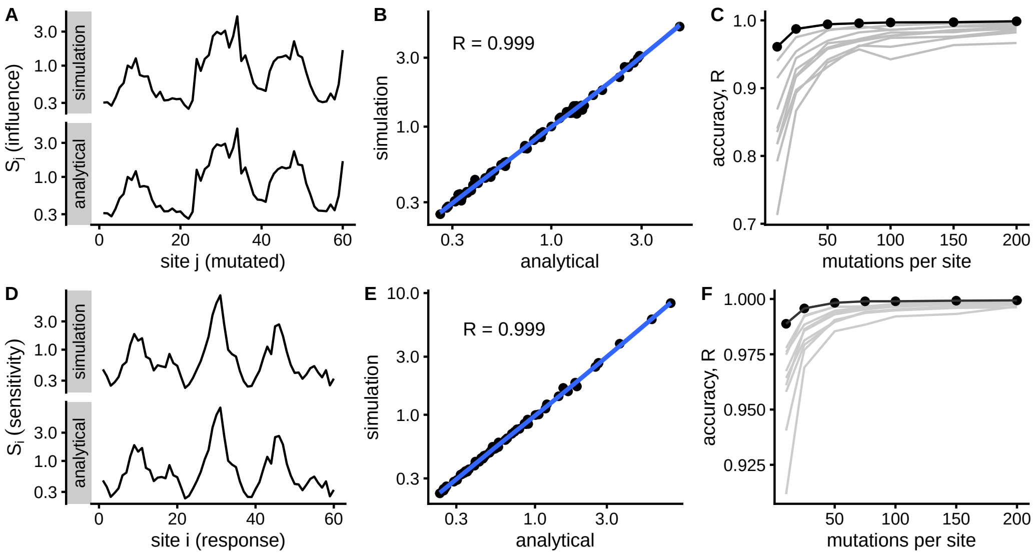

**Compensation matrix, d1ntxa**

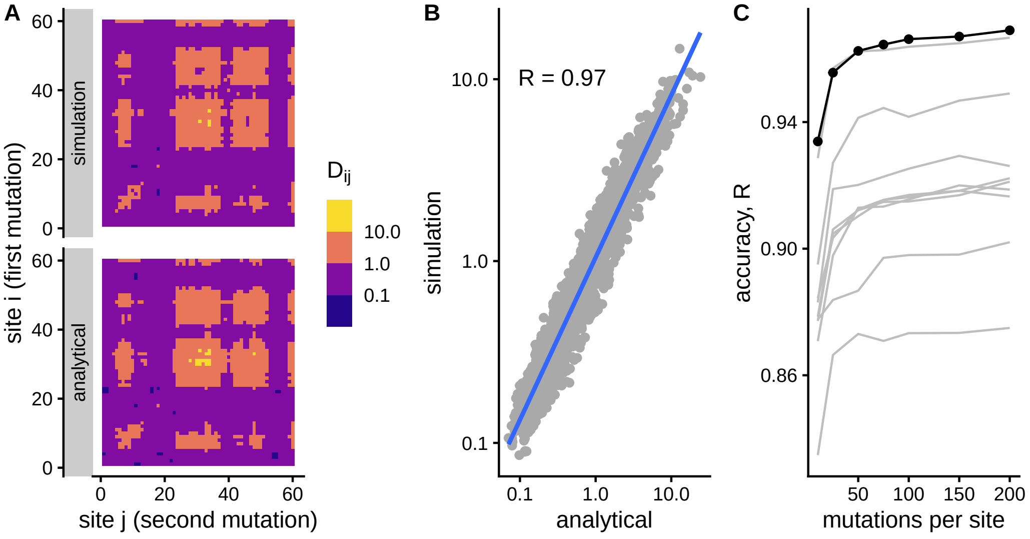


**Compensation profiles, d1ntxa**

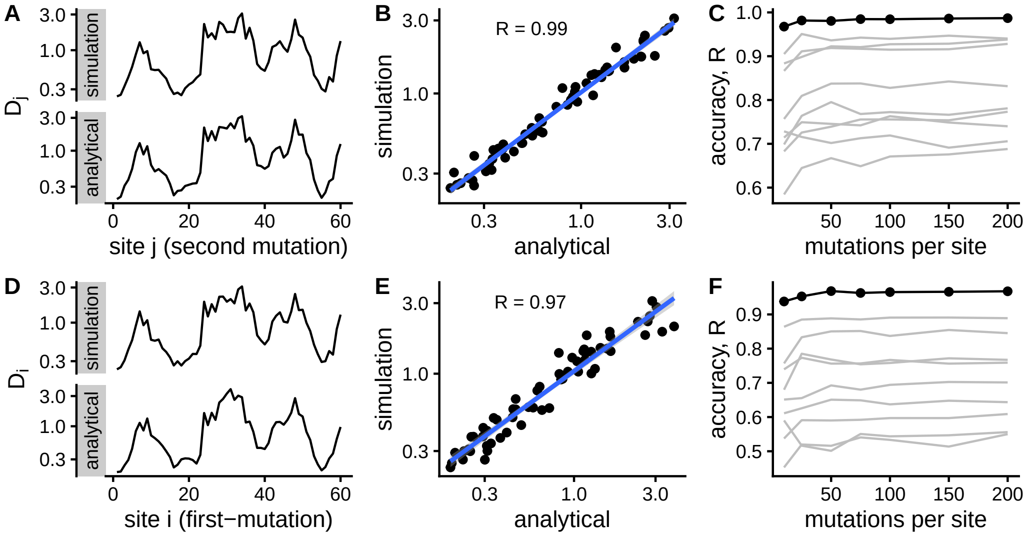

#### Sensitivity matrix, d1fxla2

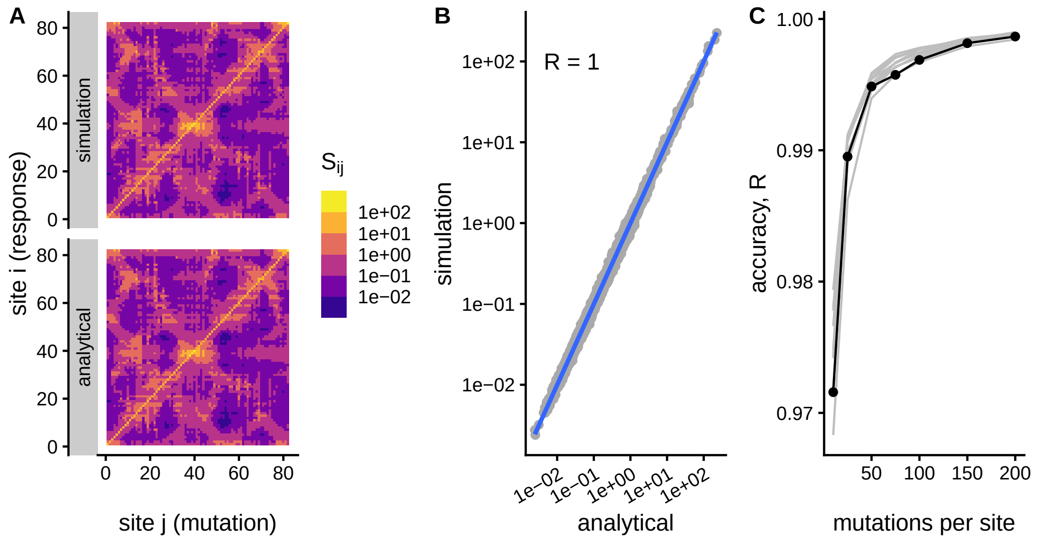


**Influence and sensitivity profiles, d1fxla2**

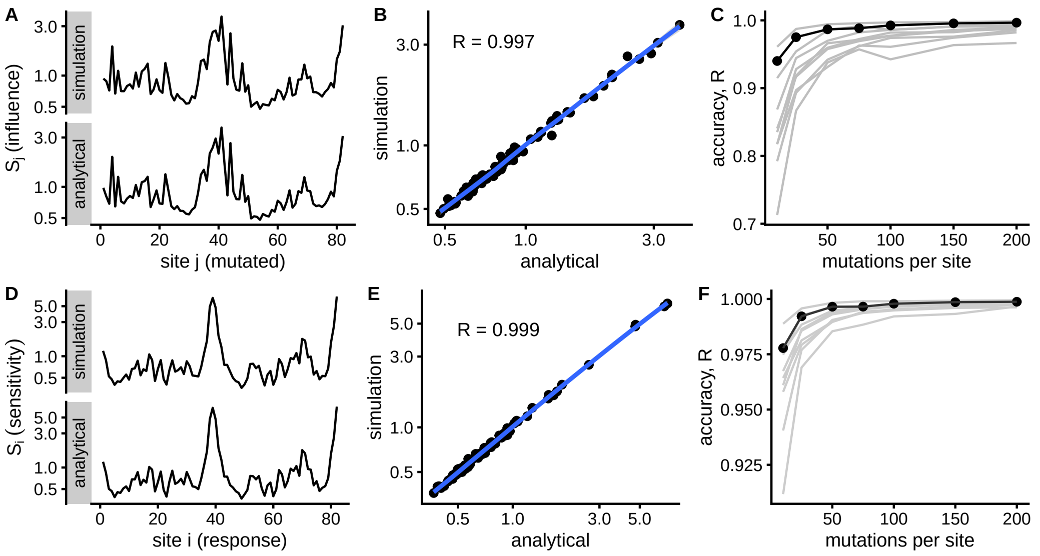

#### Compensation matrix, d1fxla2

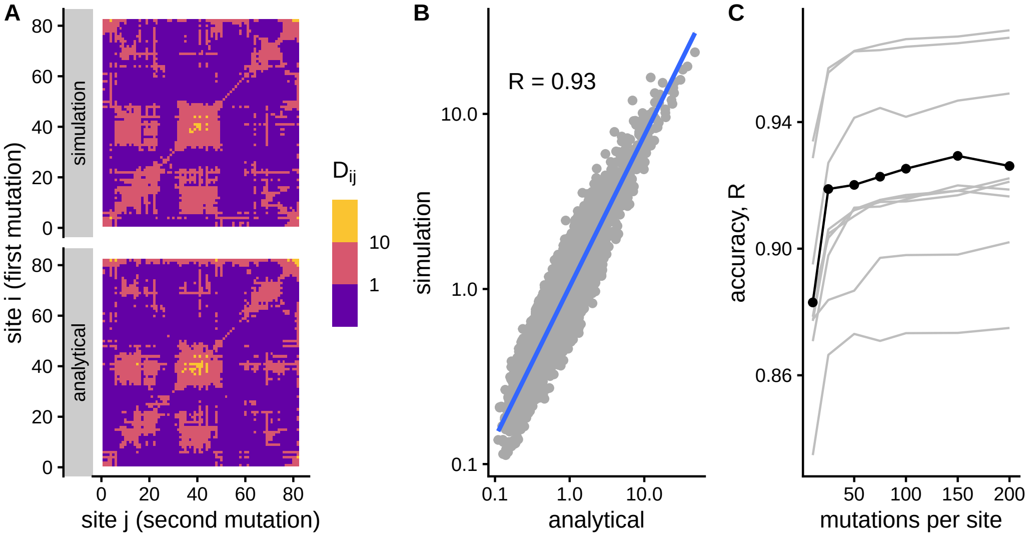


#### Compensation profiles, d1fxla2

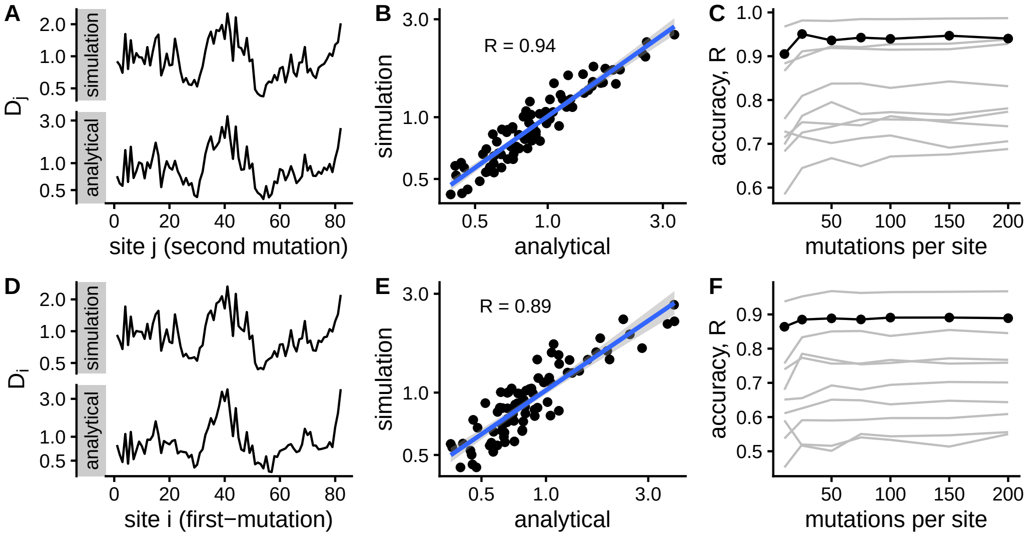

**Sensitivity matrix, d1bxva**

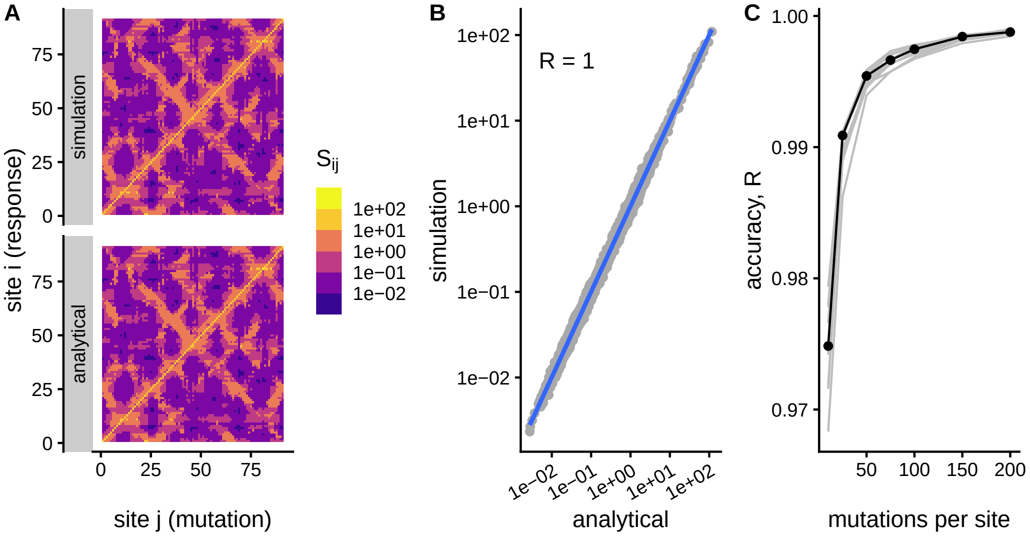


**Influence and sensitivity profiles, d1bxva**

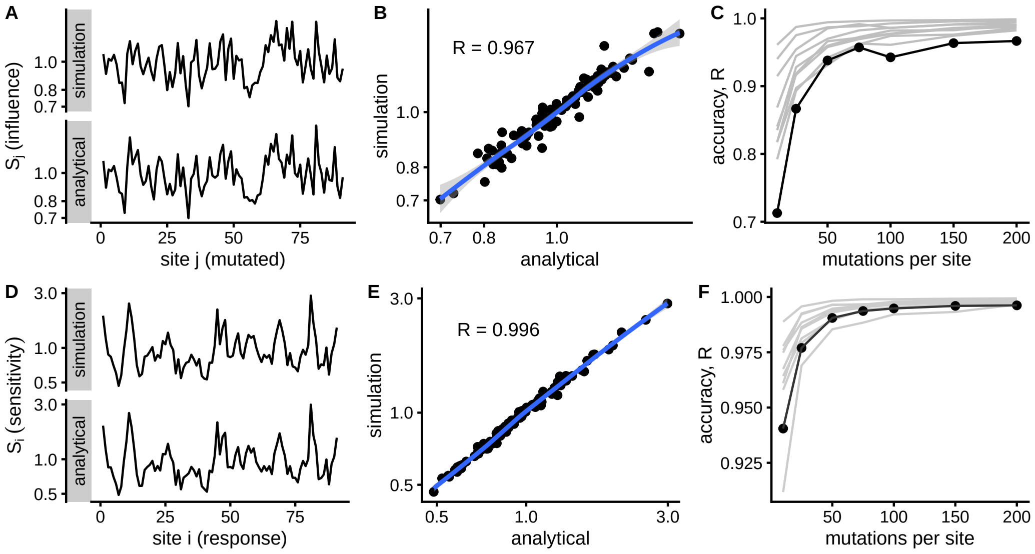

#### Compensation matrix, d1bxva

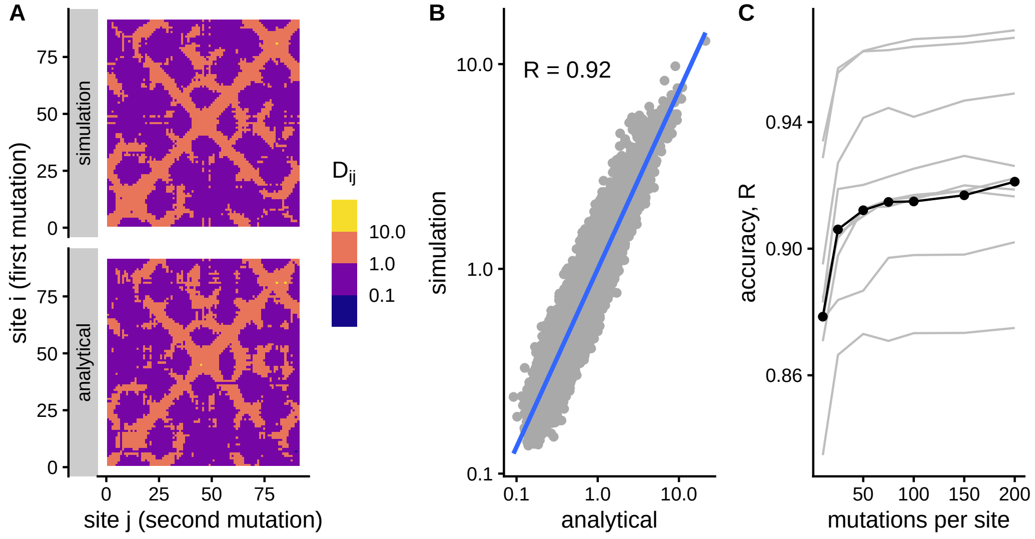


#### Compensation profiles, d1bxva

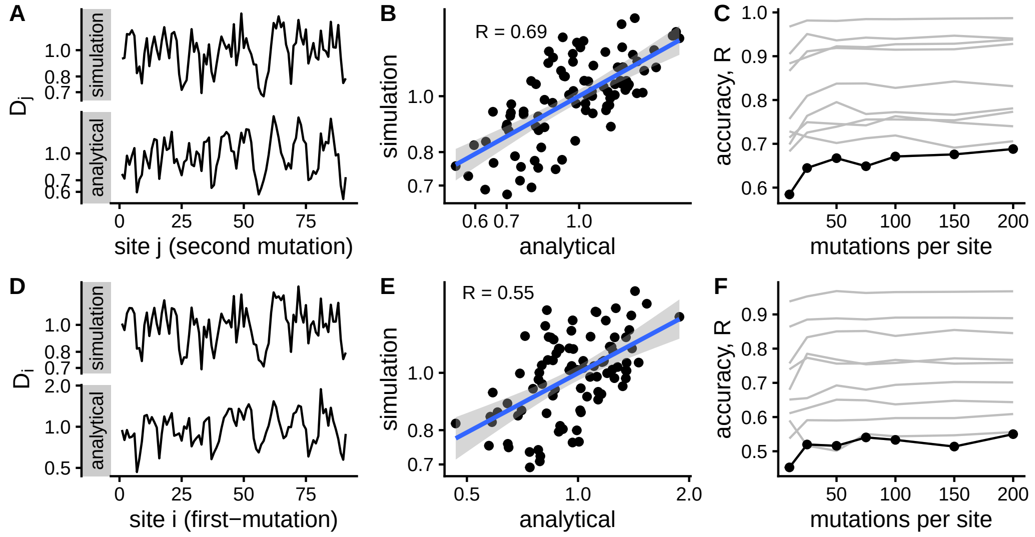

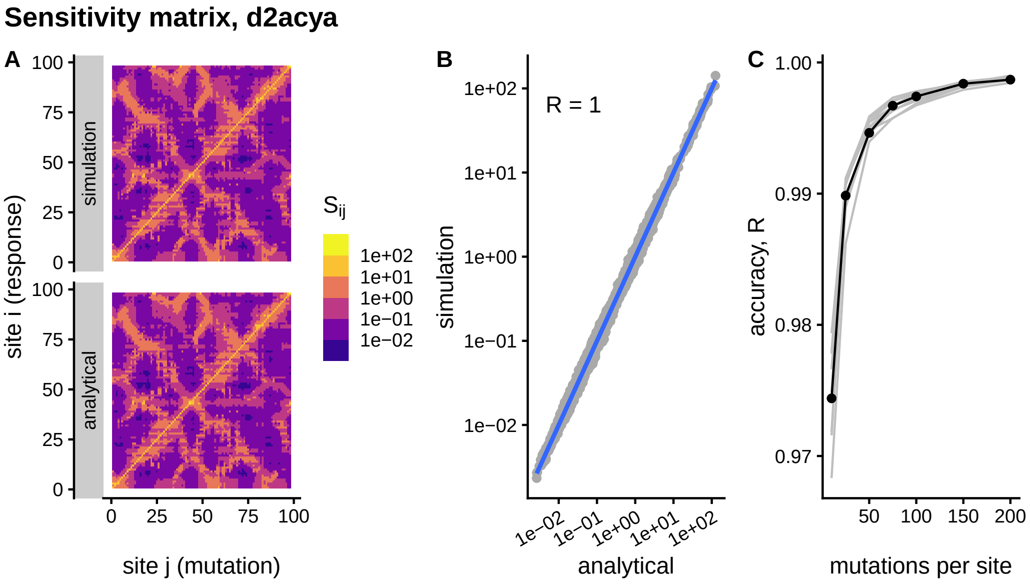
Sensitivity matrix, d2acya
A
B
C
simulation
analytical
site i (response)
site j (mutation)
S_ij
1e+02
1e+01
1e+00
1e−01
1e−02
R = 1
simulation
analytical
accuracy, R
mutations per site

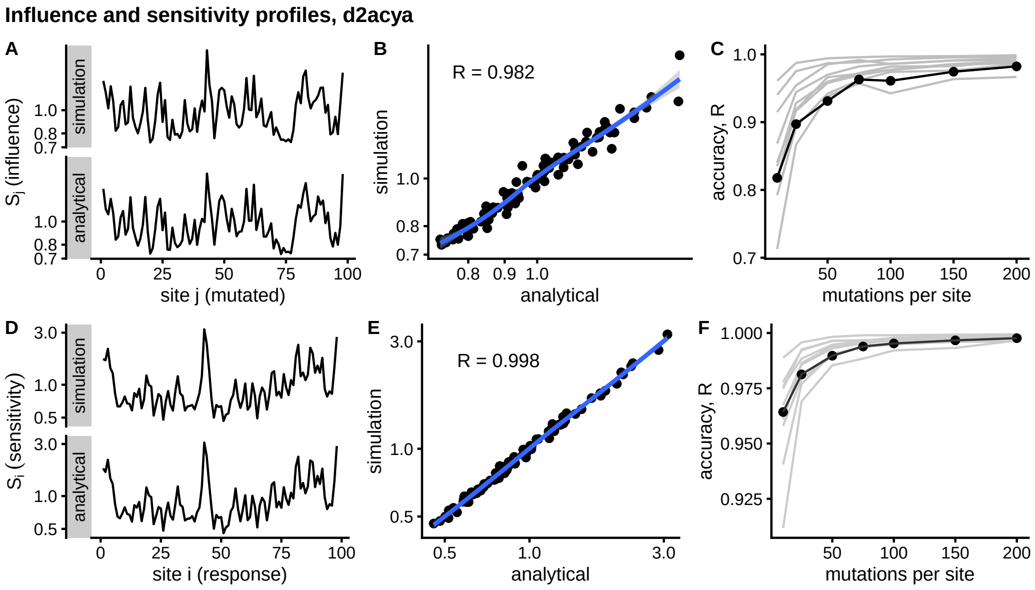
Influence and sensitivity profiles, d2acya
A
B
C
D
E
F
simulation
analytical
S_j (influence)
site j (mutated)
R = 0.982
R = 0.998
accuracy, R
mutations per site
S_i (sensitivity)
site i (response)

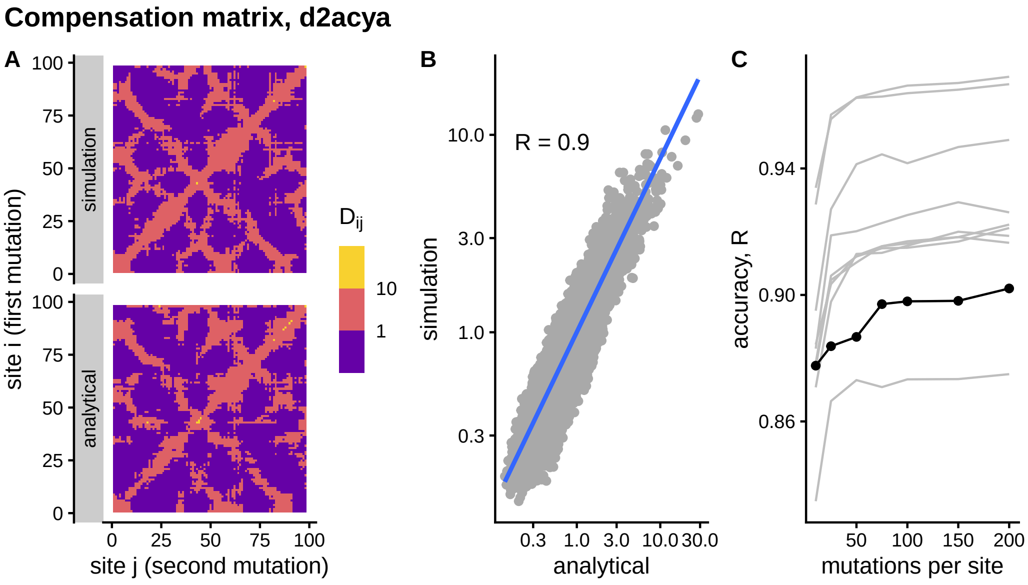
Compensation matrix, d2acya
A
B
C
simulation
analytical
site i (first mutation)
site j (second mutation)
Dij
10
1
R = 0.9
simulation
analytical
accuracy, R
mutations per site

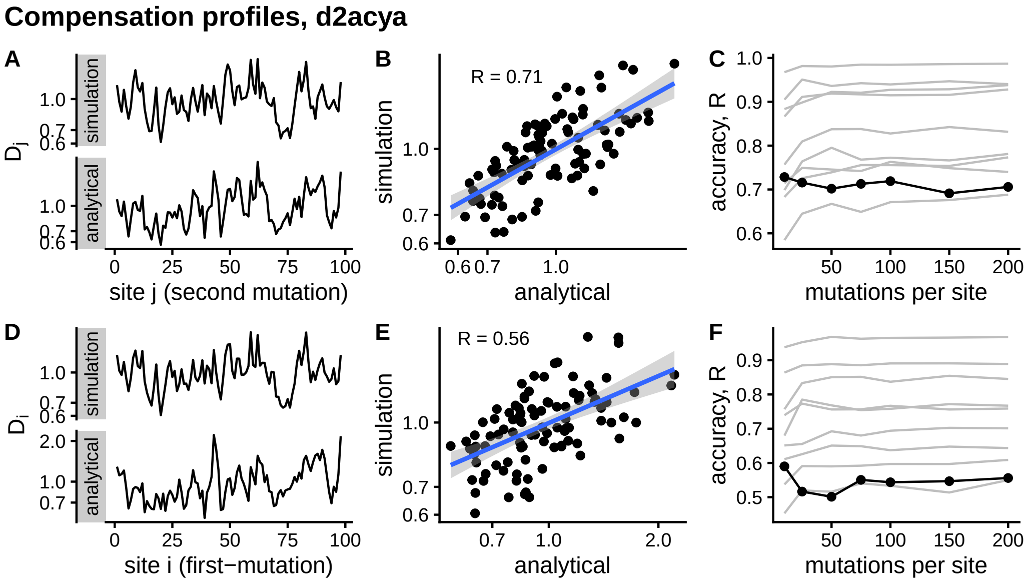
Compensation profiles, d2acya
A
B
C
D
E
F
simulation
analytical
site j (second mutation)
site i (first−mutation)
R = 0.71
R = 0.56
accuracy, R
mutations per site

**Sensitivity matrix, d1hmta**

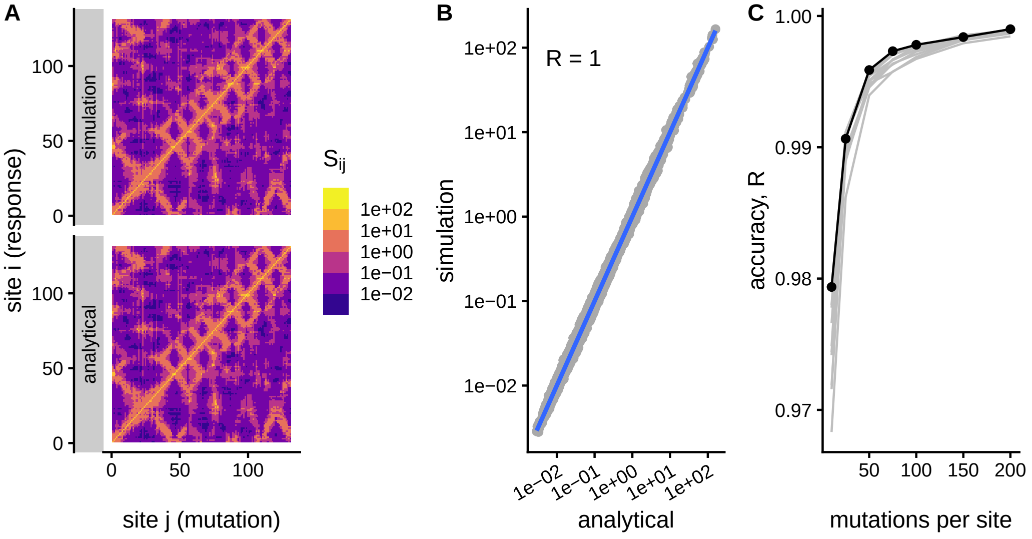


**Influence and sensitivity profiles, d1hmta**

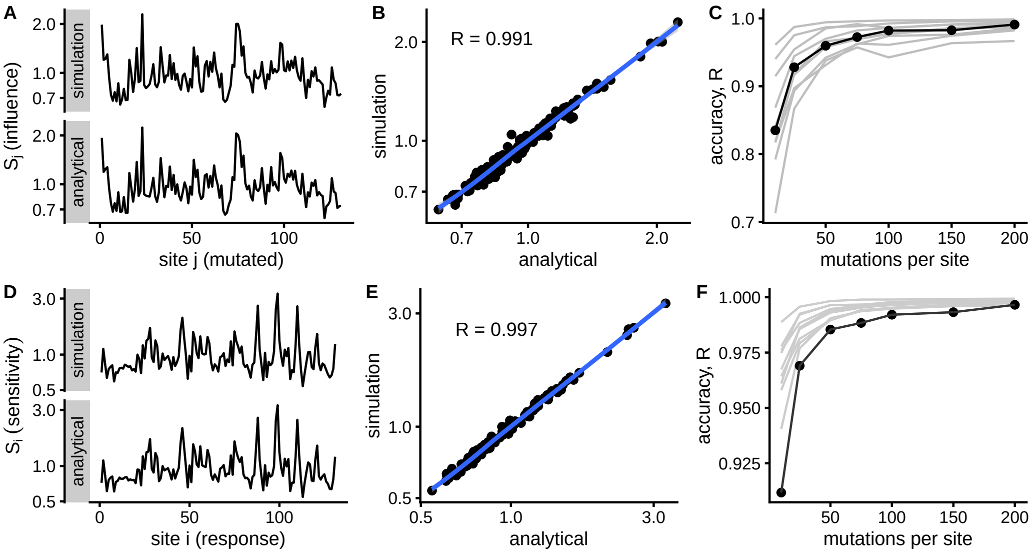

**Compensation matrix, d1hmta**

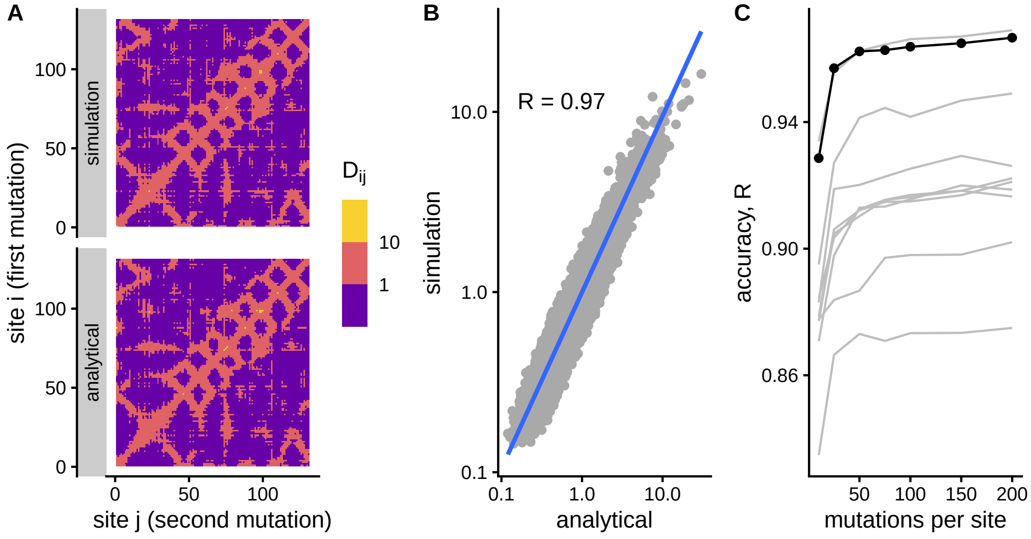


**Compensation profiles, d1hmta**

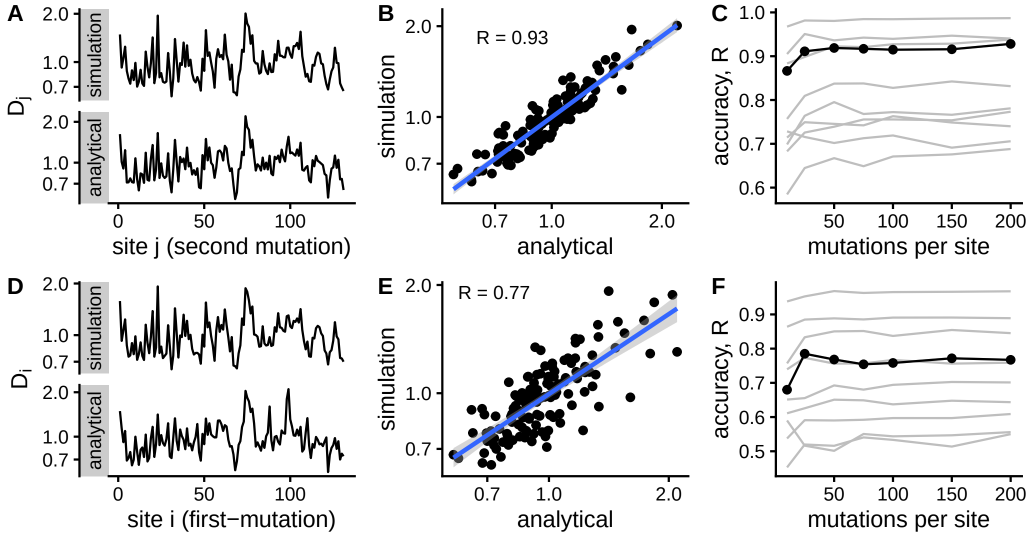

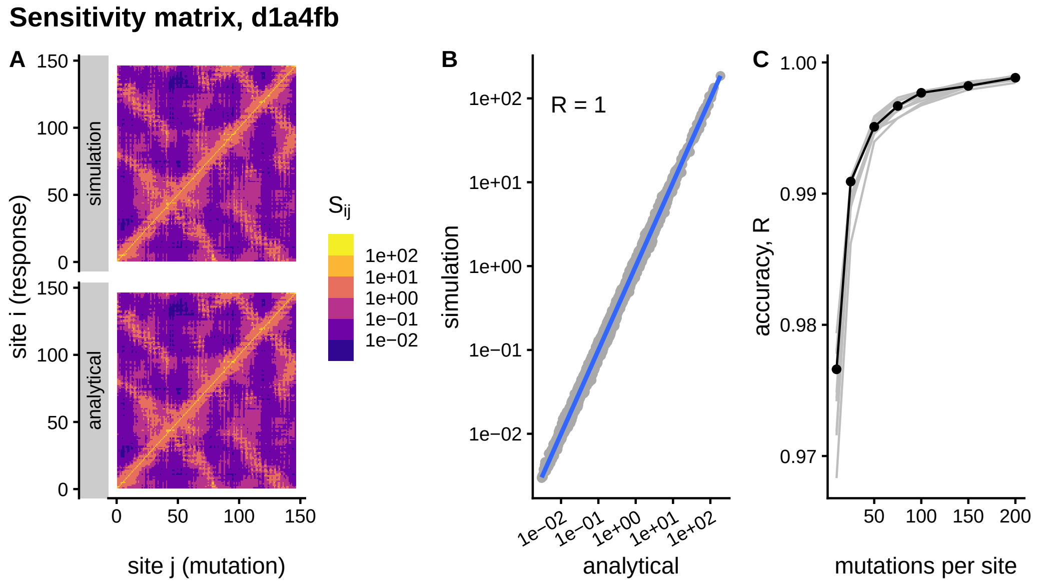
Sensitivity matrix, d1a4fb
A
B
C
simulation
analytical
site i (response)
site j (mutation)
$S_{ij}$
1e+02
1e+01
1e+00
1e−01
1e−02
R = 1
simulation
analytical
accuracy, R
mutations per site

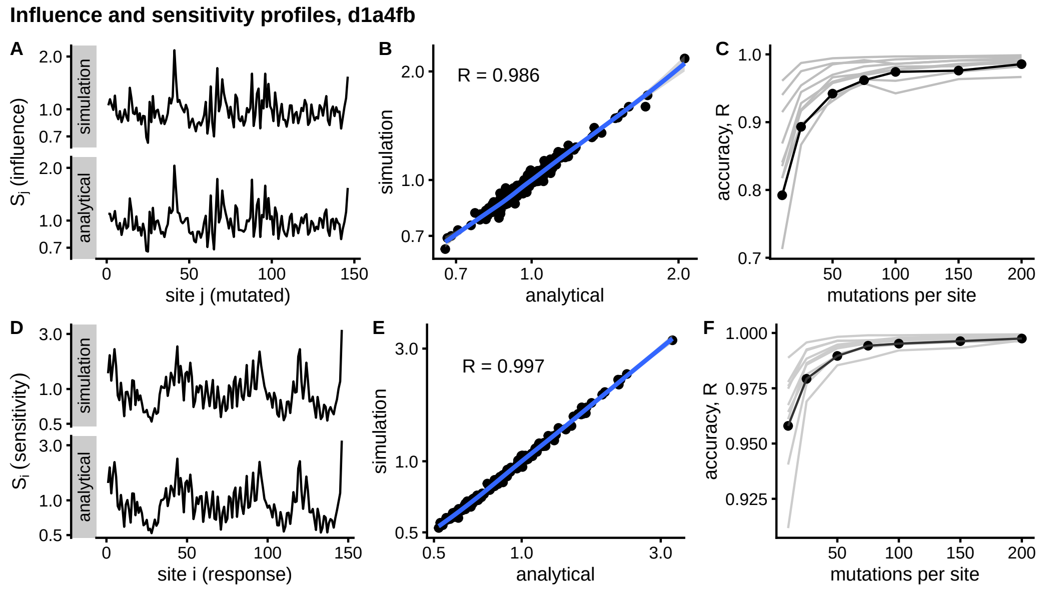
Influence and sensitivity profiles, d1a4fb
A
B
C
D
E
F
$S_j$ (influence)
$S_i$ (sensitivity)
simulation
analytical
site j (mutated)
site i (response)
R = 0.986
R = 0.997
accuracy, R
mutations per site

**Compensation matrix, d1a4fb**

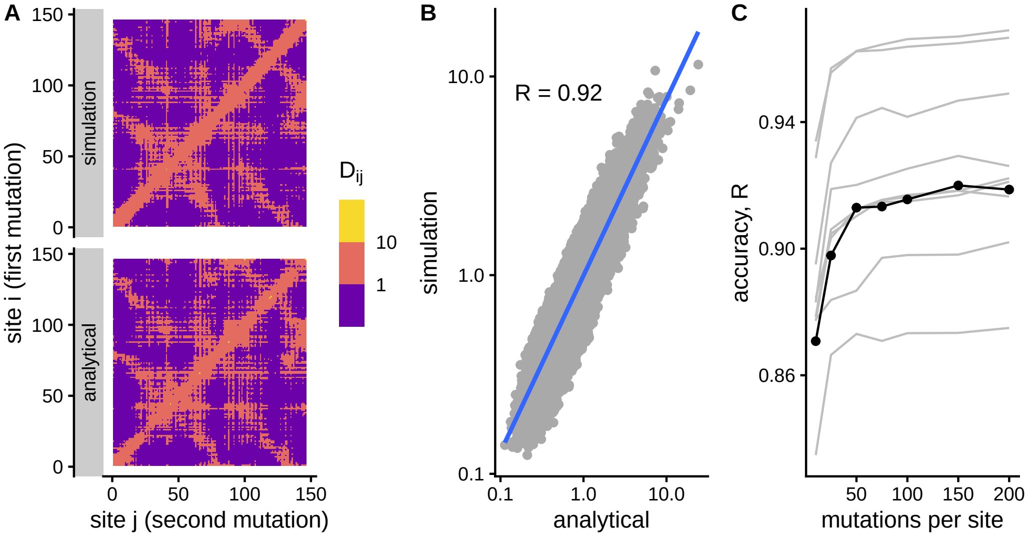


**Compensation profiles, d1a4fb**

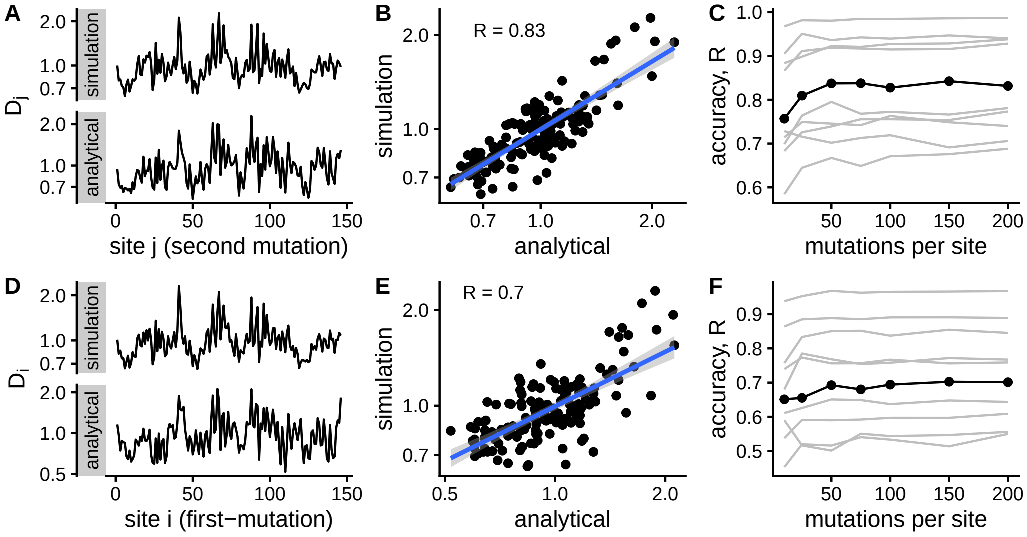

**Sensitivity matrix, d1mcta**

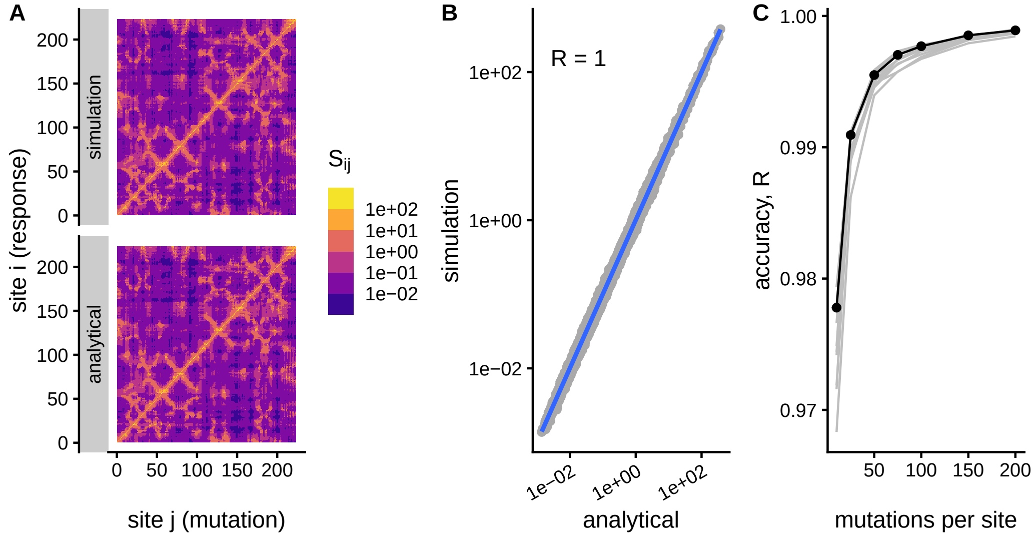


**Influence and sensitivity profiles, d1mcta**

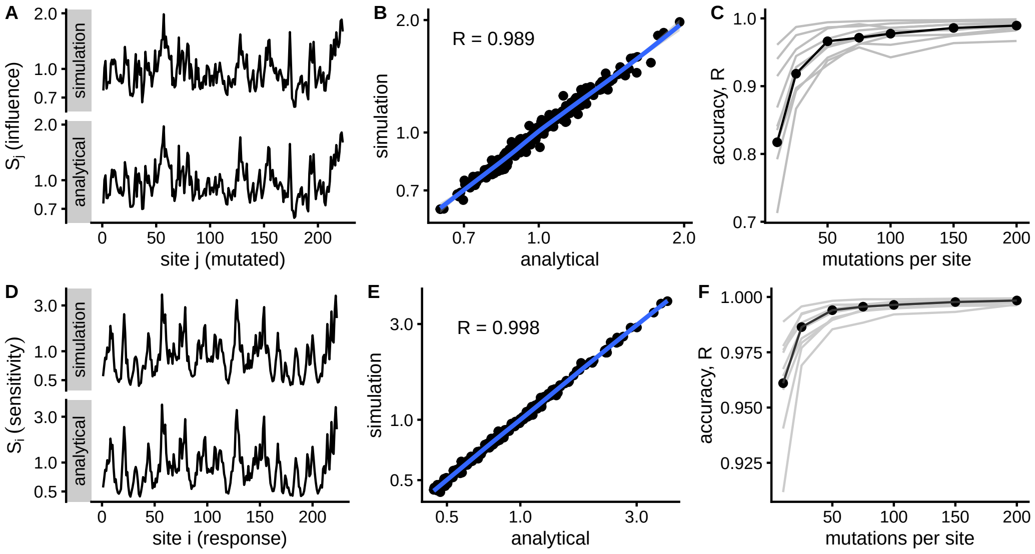

**Compensation matrix, d1mcta**

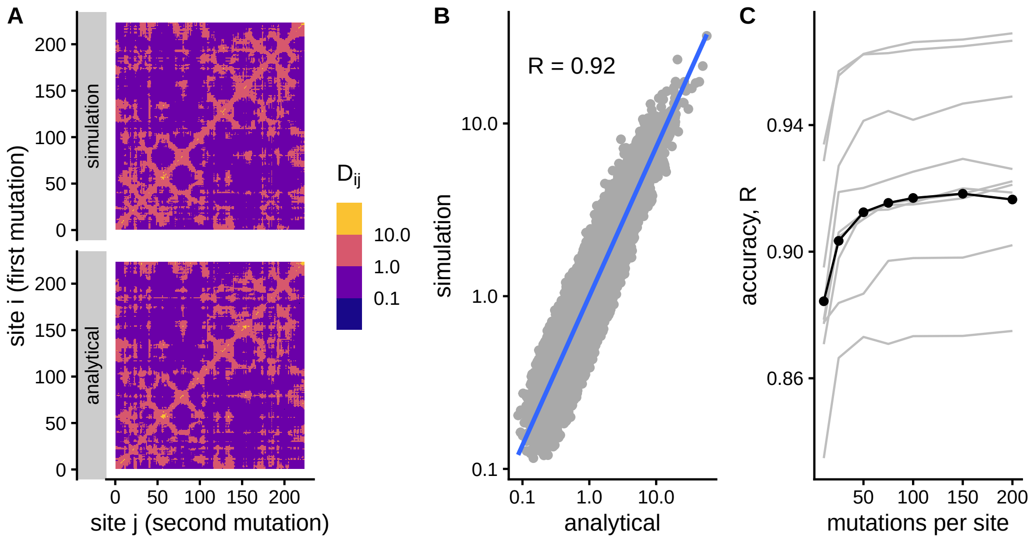


**Compensation profiles, d1mcta**

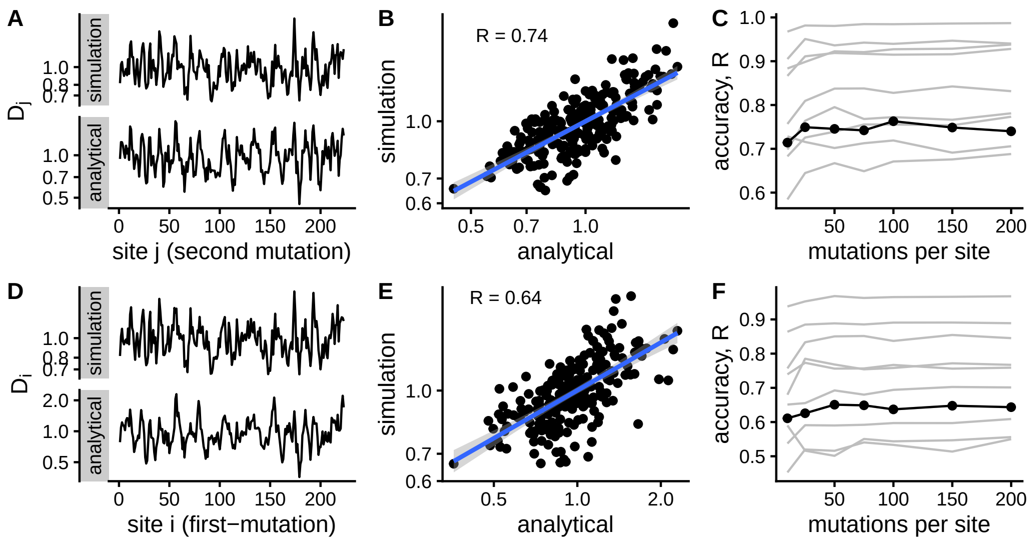

**Sensitivity matrix, d2l8ma**

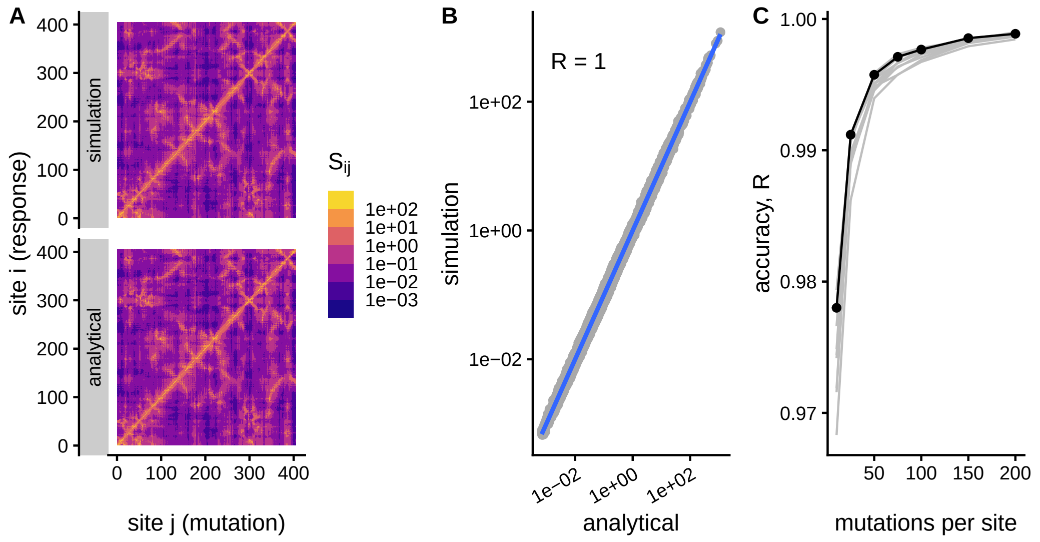


**Influence and sensitivity profiles, d2l8ma**

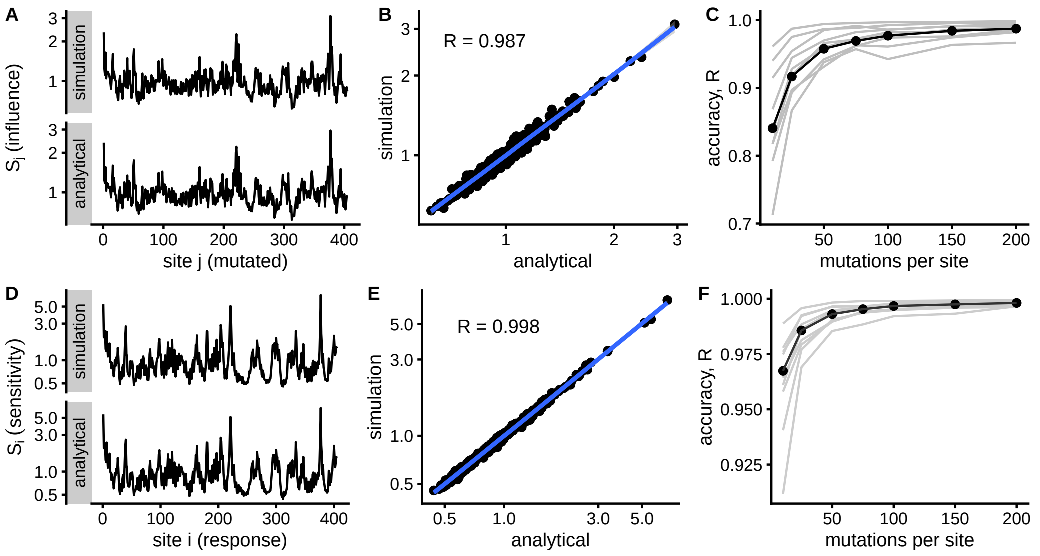

**Compensation matrix, d2l8ma**

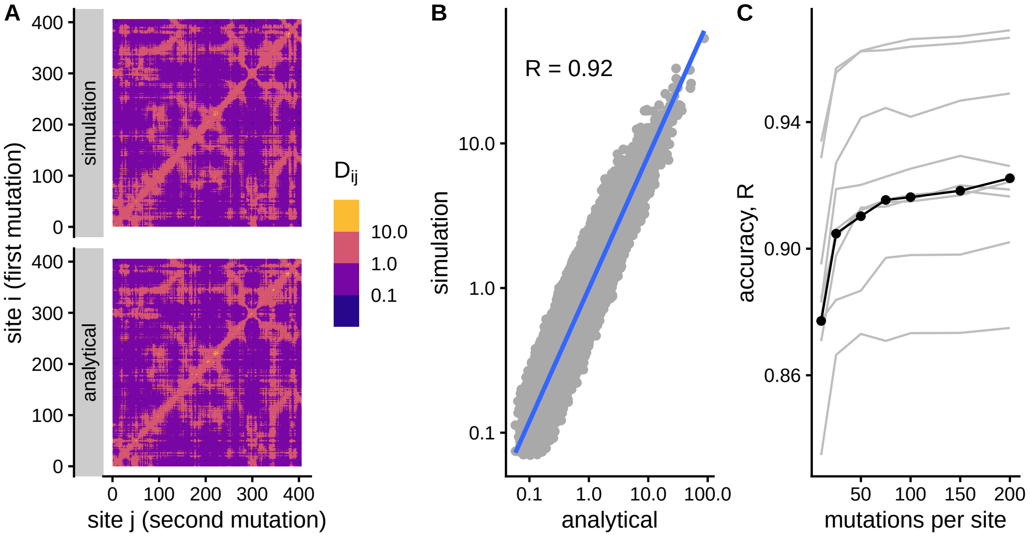


**Compensation profiles, d2l8ma**

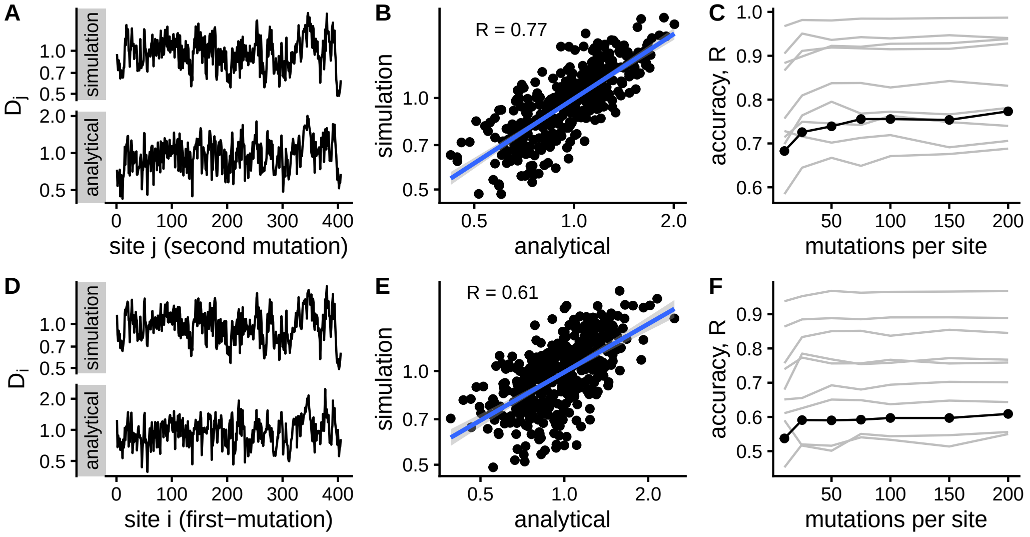
